# Cleland immunoblotting unmasks unexpected cysteine redox proteoforms

**DOI:** 10.1101/2024.09.18.613741

**Authors:** James N. Cobley, Anna Noble, Matthew Guille

## Abstract

Cysteine redox proteoforms are virtually unstudied because they are extremely difficult to detect. They are difficult to detect by immunoblotting when the polyethylene glycol (PEG)-payloads used to mobility-shift proteoforms into distinct bands block antibody binding. Here, we synthesised a novel compound to reversibly crosslink the PEG-payloads to oxidised cysteines using disulfide bonds. To reductively release the PEG-payloads from mobility-shifted proteoforms, we soaked the gel in Cleland’s reagent: 1,4-dithiothreitol (DTT). Hence, Cleland immunoblotting. Cleland immunoblotting unmasked hitherto undetectable cysteine redox proteoforms. In *Xenopus* oocytes, we detected 2 cdc20-specific coordinates in an otherwise abstract space housing 1,024 theoretical proteoforms. The coordinates mapped to the fully, all 10 cysteine residues, 100%-reduced and 100%-oxidised proteoforms. The unexpected absence of partially oxidised molecules along the 10-reaction path to the 100%-oxidised proteoform implies novel biology. By providing the technological means to detect cysteine redox proteoforms, Cleland immunoblotting opens new avenues of discovery.

## 1. Introduction

Understanding redox biology depends on describing the cysteine redox state of proteins [1]. What is the cysteine redox state of a protein? The cysteine redox state of a protein molecule is a function of a finite number of cysteine redox proteoform (*i*) states [2], as mathematically defined in equation 1. As codified in equation 2, every protein molecule with a cysteine residue (P_CYS_) is a proteoform. Their formal equivalence means that proteoform *i* states define the fundamental description of protein redox biology. However, the *i* states that proteins adopt are unknown as cysteine redox proteoforms are virtually unstudied [3].

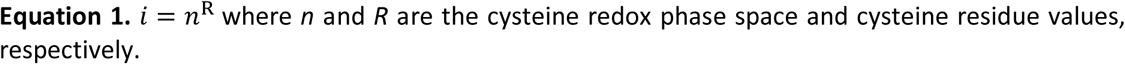

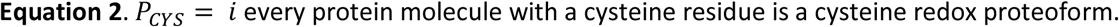

Cysteine redox proteoforms are virtually unstudied because they are extremely challenging to detect [3]. For example, the immunoblotting-based m-PEG assay struggles to detect cysteine redox proteoforms because the mobility-shifting polyethylene glycol (PEG)-payloads usually block antibody binding [4–6], resulting in the measurement of only ≈30 proteins in over 20 years [7]. Equally pressing challenges beset mass spectrometry (MS)-based methods [8]. Most proteoform information is irretrievably lost when proteins are digested into peptides in bottom-up MS [9]. Meanwhile, top-down MS struggles to match peptides to specific proteoforms, suffers from insufficient residue coverage, and is relatively insensitive [10]. The technical challenges responsible for making the *i* states that proteins do adopt unknowable rate-limit our understanding of redox biology.

Knowing the *i* states proteins adopt is fundamental for understanding the redox biology of protein function, post-translational regulation, and oxidative stress in health and disease [11]. All of the above, with all of their profound importance to virtually every research topic in the life sciences [12–14], are actuated by cysteine redox proteoforms. For example, protein function often relies on assuming specific *i* states, such as cysteine redox proteoforms with a reduced active site thiolate in the case of many enzymes [15]. Redox regulation exploits these dependencies to post-translationally control protein function [16], converting one proteoform into another. Networked cysteine redox proteoform system states contribute to oxidative stress [17]. And yet, the cysteine redox states of proteins, as codified by equation 3, are undefined. Take a protein with 5 cysteines. Each molecule must adopt one of 32 *i* states. If the population of molecules were 20%-oxidised, then the redox state could be a product of up to 6 percent-oxidised proteoform redox grades per equation 4:

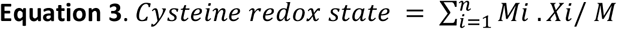

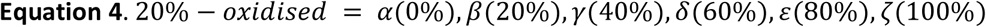

Computing equation 4 using the <u>Cys_%ox_cal.py</u> script (supplementary data file 1), yielded 30 possible ways to solve a 20%-oxidised cysteine redox state for a population (*M*) of just 10 molecules. Computing larger *M* values is challenging owing to the exponential increase in the number of possible solutions as a function of *M*. The sheer number of potential combinatorial solutions means the nature of the proteoforms that define the redox states of proteins cannot be deduced from a set of potential solutions. Their states are essentially unknowable without novel technologies for making empirical measurements. Hence, the ability to make empirical measurements could open new avenues of discovery by defining the *i* states that actuate protein biology, redox regulation, and oxidative stress.

As a powerful technology [18], immunoblotting provides an ideal methodological basis for measuring cysteine redox proteoforms. Overcoming the longstanding PEG-payload problem could finally unmasking the elusive proteoforms that define the redox states of proteins. Such a breakthrough would establish a foundational methodological platform for advancing redox biology. However, unlocking these insights hinges on solving the PEG-payload problem [7].

## 2. Results and discussion

### 2.1. A novel solution to the PEG-payload problem

To solve the PEG-payload problem, we formalised a novel concept where the PEG (*P*) and antibody-antigen binding (*A*) states are encoded in binary where 0 and 1 are “OFF” and “ON”, respectively. The only condition is that: the protein must be detectable by immunoblotting when *P* = *0*. The PEG-free state. When this condition is met, we proposition that the probability of successfully (*P_success_*) detecting cysteine redox proteoforms is 1, which can be mathematically expressed as:

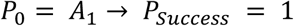

If they are all are in a uniform *P_0_* state, then the antibody will detect all of the antigens regardless of their position on the membrane. The uniform *P_0_* state means the PEG-payloads cannot block antibody binding. Hence, cysteine redox proteoforms will invariably be detected. To illustrate, imagine the catalytic subunit of the protein phosphatase PP2A (<u>PPP2CA</u>), use the <u>Cys_PEG_PDB.py</u> script to generate the PEG-modified for a protein of your choosing, in 3 method-dependent states:

1. **Western**. In a Western blot, the antibody binds to a linear version of the PPP2CA structure displayed in Figure 1, in which all ten cysteine residues are in the *P* = 0 state. Hence, a band at ≈37 kDa can be detected. However, no proteoforms can be resolved without the PEG-payloads (i.e., *P_0_* = *i_0_*). **Outcome**: *P_0_* = *A_1_*.
2. **m-PEG.** In the m-PEG assay, the antibody must bind a linear version of the PPP2CA structure displayed in Figure 1, in which all ten cysteine residues are in the *P* = 1 state. In the *P_1_* state, PPP2CA will be mobility-shifted by ≈20 kDa. However, no proteoforms can be detected as the PEG-payloads block antibody binding, as evidenced for PPP2CA [19]. **Outcome**: *P_1_* = *A_0_*.
3. **Proposition**: In our formal proposition, PPP2CA is in the *P_1_* state during electrophoresis and then converted to the *P_0_* state before the membrane transfer step. Hence, the antibody can bind to the mobility-shifted cysteine redox proteoforms. **Outcome**: *P_1_* → *P_0_*= *A_1_*.

**Figure 1.**
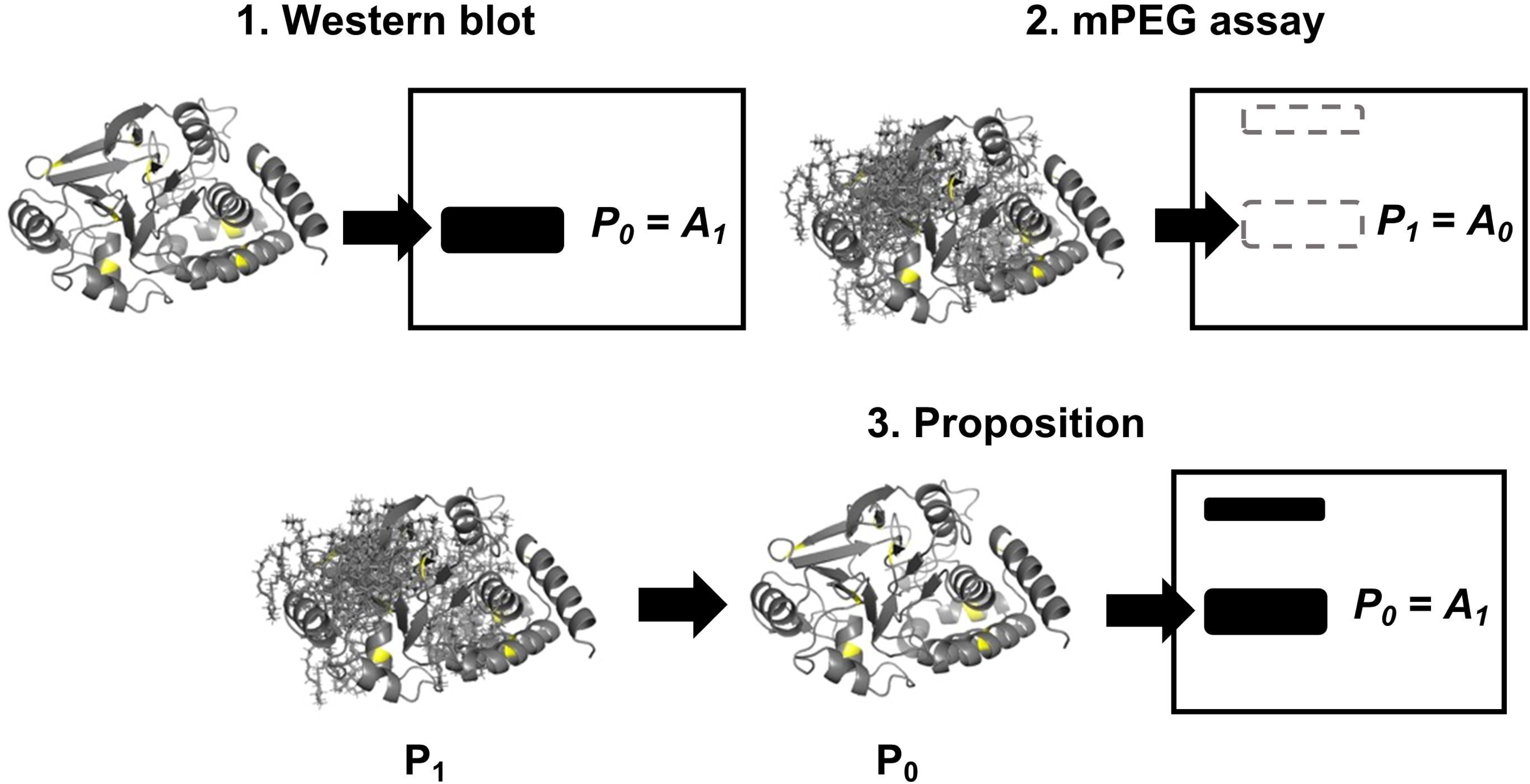
Structures of human <u>PPP2CA</u> in three method-dependent states. In Western blotting, the protein is free of the PEG-payloads (*P* = 0) enabling the antibody to detect the antigen, resulting in the appearance of a band. However, without the PEG-payloads no cysteine redox proteoforms are detected. In the m-PEG assay, cysteine redox proteoforms are mobility-shifted by the PEG-payloads (*P* = 1). However, the PEG-payloads block antibody binding. In the formal proposition, the PEG-payloads are present (*P* = 1) to mobility-shift cysteine redox proteoforms before being removed (*P* = 0). The uniform *P_0_* state should enable all of the target protein antigens to be detected by the antibody regardless of their position on the membrane. The structure for human PPP2CA was downloaded from <u>Alphafold</u>.

The only difference between a Western blot and our formal proposition is where the antigen is on the membrane. Antibody binding is independent of membrane position. For example, in a dot blot, one could spot the antigen in any of the membrane coordinates and still detect it, provided the epitope is displayed. The location of the signal is a function of position, whereas obtaining a signal is a function of antibody binding. This critical distinction underscores that the key factor for successful detection is the presence of the epitope in a state that can be recognised by the antibody.

In sum, we defined the conceptual basis of a novel immunoblot-based technology for detecting cysteine redox proteoforms by elaborating detective arguments that should produce a successful outcome when they are satisfied.

### 2.2. Cleland Immunoblotting technology

We have defined deductive arguments that should logically guarantee the detection of all cysteine redox proteoforms regardless of their position on the membrane. Under the condition of a variable *P_1_* state, cysteine redox proteoforms will be mobility-shifted in an oxidation-dependent manner. Hence, the PEG-payloads must be present during SDS-PAGE. After SDS-PAGE, the PEG-payloads must be removed to achieve a uniform *P_0_* state. Under the condition of a uniform *P_0_* state, the detection is oxidation-independent: the antibody should be able to bind all antigens regardless of their position on the membrane. Realising this concept required the chemical means to flick the P-state switch between the electrical SDS-PAGE and membrane transfer immunoblot modules. Inspired by the very proteoforms we sought to detect [2], we designed a novel cysteine redox state change sequence for flicking the P-state switch:

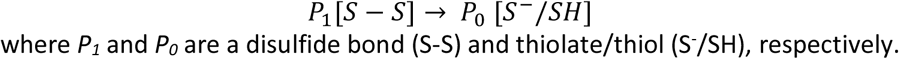

where *P_1_*and *P_0_* are a disulfide bond (S-S) and thiolate/thiol (S^-^/SH), respectively.

To realise this redox sequence, we designed a new compound called <u>2</u>-pyridyldithiol-<u>P</u>EG_47_-<u>b</u>iotin (2PB) using “Lego-like” building blocks (supplementary Figure 1). In a simple 1-pot and enzyme-free reaction, 2-pyridyldithiol-PEG_36_-*N*-hydroxysuccinimide (NHS) and amine(NH_2_)-PEG_11_-biotin are directionally united via an amide bond per reaction 1.

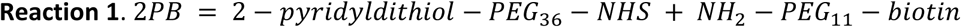

The 2-pyridyldithiol group should selectively label cysteine residues (C) via a disulfide bond exchange reaction that releases pyridine-2-thione (PS) from 2PB. The stable 2PS leaving group enhances the kinetics of the disulfide exchange reaction by lowering the activation energy. An added bonus: the 2PS thioketone tautomer cannot participate in further redox reactions.

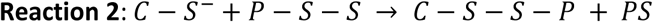

The C-S-S-P disulfide bond mobility-shifts cysteine redox proteoforms by their cysteine oxidation integer in non-reducing SDS-PAGE. After SDS-PAGE, the PEG-payloads are released by reducing the C-S-S-P disulfide bond in-gel using Cleland’s reagent: DTT [20]. Ergo, Cleland immunoblotting (Figure 2). Cleland immunoblotting can measure cysteine redox proteoforms, provided (a) protein can be detected by immunoblotting and (b) the concentration of antigen in a mobility-shifted band exceeds the limit of detection.

**Figure 2.**
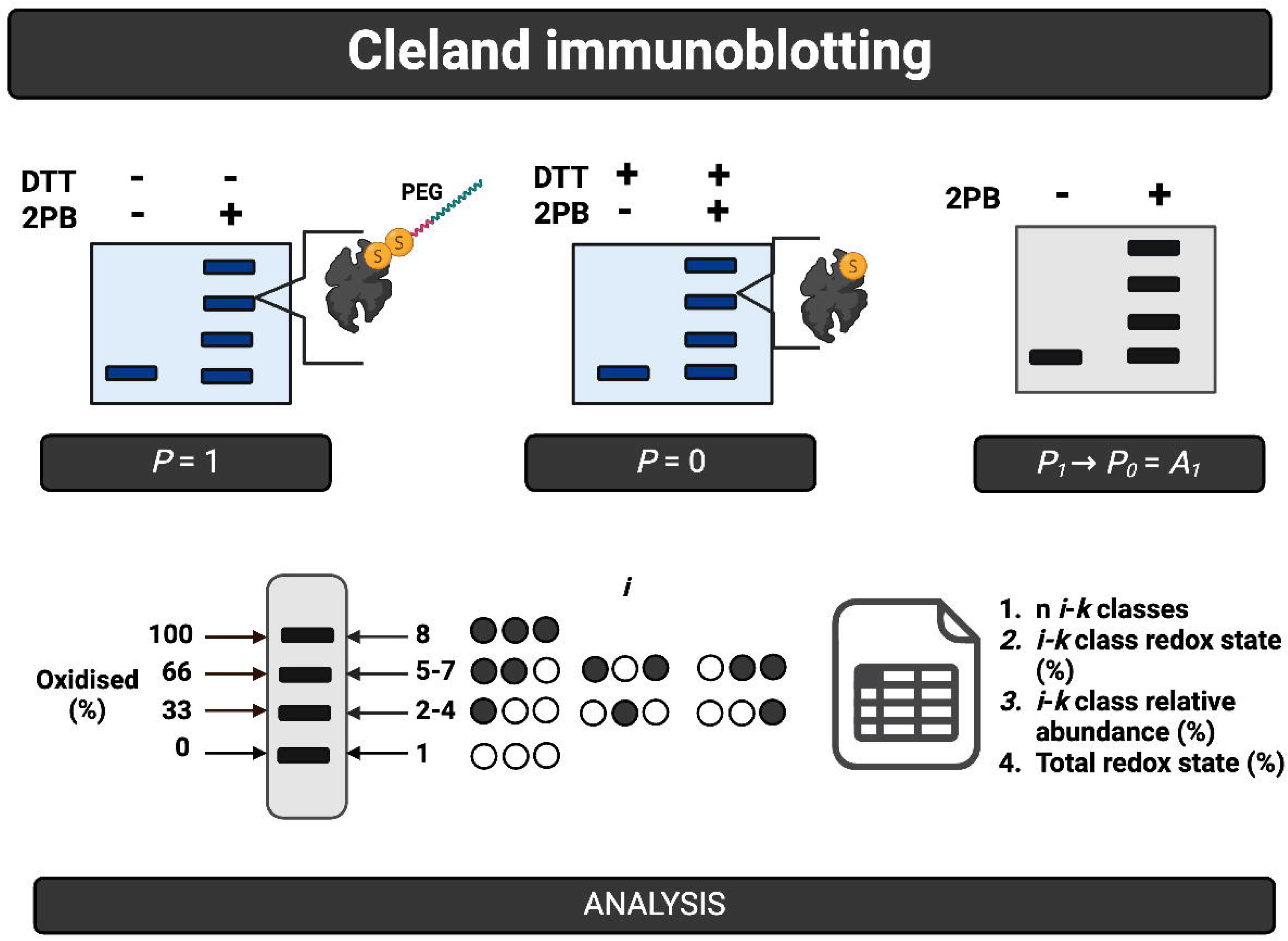
Cleland immunoblotting technology. After oxidised cysteine residues are labelled with 2PB (see main text), cysteine redox proteoforms (*i*) are mobility-shifted by their cysteine oxidation integer (*k*) in SDS-PAGE due to the presence of the PEG-payloads (*P* = 1). The PEG-payloads are then reductively released by soaking the gel in DTT—Cleland’s reagent. Hence, Cleland immunoblotting. Placing all of the target antigens, regardless of their position on the membrane, in the uniform *P* = 0 state, enables antibody binding (*A* = 1). Hence, *P_1_ →P_0_*= *A_1_*. As illustrated for a protein with 3 cysteines (*r* = 3), the 8 theoretical cysteine redox proteoforms are stratified into 4 percentage redox grade bands. Each band is defined by a specific *k* value (e.g., 0%-oxidised = *k* 0). The distribution of cysteine redox proteoforms into *k*-defined bands is determined by binomial theorem. Overall, Cleland immunoblotting can identify and quantify cysteine redox proteoforms.

The maximum number of mobility-shifted bands (*nB*) that can be identified by Cleland immunoblotting is a function of the cysteine residue integer (*r*) plus 1 per equation 5. For example, a protein with 3 cysteine residues, such as <u>GAPDH</u>, can form 4 bands. The percentage redox grade increment (*g*) between the bands is *r* divided by 100 per equation 6. For example, *g* values for GAPDH are: 0, 33.3, 66.6, and 100%-oxidised (or 100, 66.6, 33.3, and 0-%-reduced). The number of unique cysteine redox proteoforms (*i*) in each cysteine oxidation integer specified (*k*) percentage redox graded band (%) is given by binomial theorem (equation 7). The binomial theorem can be geometrically visualised using Pascal’s triangle (Figure 3). For example, the geometrically-defined band structure for GAPDH is 1:3:3:1 for the 0, 33.3, 66.6, and 100%-oxidised bands, respectively.

**Figure 3.**
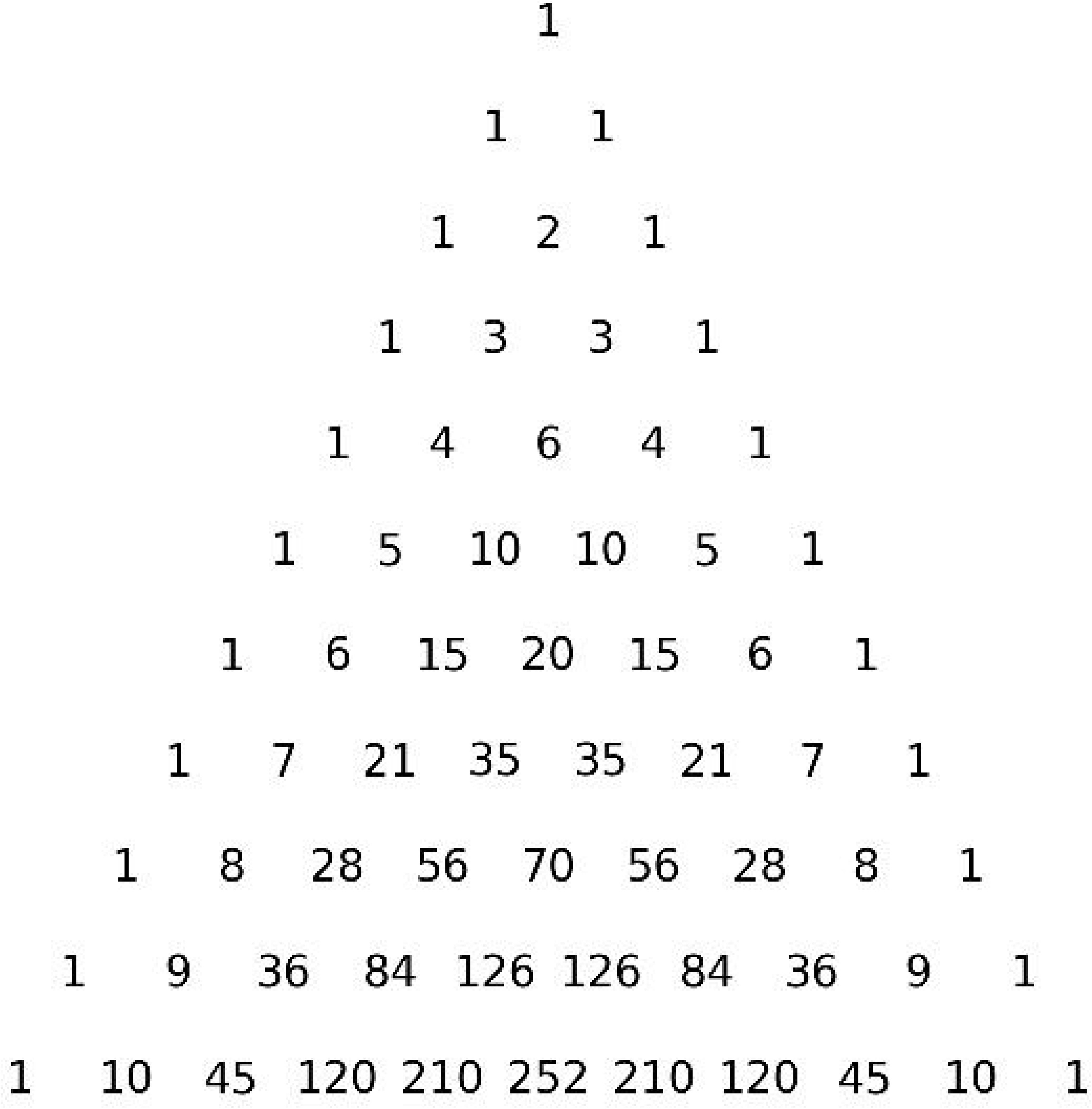
Pascal’s triangle. Geometrically illustrated *k* value defined *i* distributions per binomial theorem. The triangle contains 10 series ordered by the cysteine residue integer (*r*). For example, series 1 contains 2 values mapping to the binary proteoforms for a protein with 1 cysteine (1,1). Hence, each band would contain one proteoform. The values displayed map to the proteoform structure of the band, such that *r* = 3 would be 1,3,3,1 for the 0,33.3, 66.6. and 100%-oxidised bands, respectively. Per the classification system (Table 1), the first and last value for each series would represent a class 1 assignment whereas all the others would be class 2B. For example, class 1 for the 0 and 100%-oxidised bands for series 3 and class 2B for the 33.3 and 66.6%-oxidised bands. The image was created using the <u>Pascal.py</u> script.

**Table 1.** A 5-level cysteine redox proteoform identification-based classification system. The system objectively assigns the observed cysteine redox proteoforms to specific level stratified classes based on the number and or type of positive identifying features.

| Class | Identifications |  |  |  |
| --- | --- | --- | --- | --- |
|  | Redox grade (%) | Cys-OX/RED | Residues | Protein |
| 1 | ✓ | ✓ | ✓ | ✓ |
| 2A | X | ✓ | ✓ | ✓ |
| 2B | ✓ | ✓ | X | ✓ |
| 3 | X | ✓ | X | ✓ |
| Unclassified | X | X | X | ✓ |

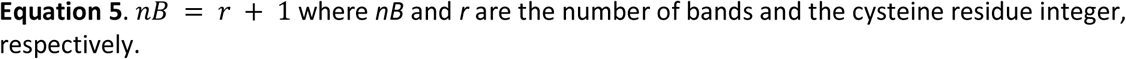

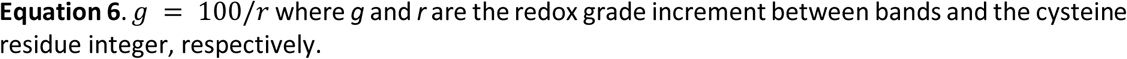

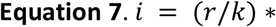

Geometric band structures inspired a cysteine redox proteoform classification system, which complements a broader rubric for classifying proteoforms [21], for clearly defining what cysteine redox proteoform technologies can identify (Table 1). In class 1, unique cysteine redox proteoforms can be unequivocally assigned. For example, there is only one way for any protein to be fully reduced or oxidised, these “binary” proteoforms can be unambiguously identified. In class 2A, the cysteine residues involved are known, but the percentage redox grade is ambiguous. In class 2B, the percentage redox grade is known, but the residues involved are unknown. For example, if one detected the 10%-oxidised band for a protein with 10 cysteines, the band could house between 1 and 10 unique cysteine redox proteoforms, leaving the oxidised residue unidentified. Class 3 assignments occur when it is only known that the protein is reduced, oxidised, or a combination thereof.

As illustrated in Figure 2 and 3, Cleland immunoblotting can identify class 1 and 2B assigned cysteine redox proteoforms. For example, in the case of GAPDH, the 0 and 100%-oxidised bands would represent a class 1 assignment. Meanwhile, the 33.3 and 66.6%-oxidised bands would represent class 2B assignments. The relative abundance of each band can be quantified in an internally normalised, within-lane, way. Hence, the overall percentage cysteine redox state of the target can be calculated using equation 3.

In sum, the Cleland immunoblotting concept defined a novel chemistry-enabled technology for measuring cysteine redox proteoforms.

### 2.3. Validating the P-state switch: The essential uniform *P_0_* condition is met

To realise Cleland immunoblotting, we validated the 2PB-based cysteine labelling procedures (Figure 4). To render reduced cysteine residues orthogonal to 2PB, they were alkylated using *N*-ethylmaleimide (NEM) [22]. NEM alkylated reduced cysteines as determined via pulse chasing with fluorescent-maleimide (F-MAL). Excess NEM was removed by a spin column, meaning no unreacted NEM can suppress 2PB labelling. As expected, Tris-(2-carboxyethyl)phosphine (TCEP) reduced oxidised cysteines. By removing the excess TCEP, the spin column step prevented it from reversing the *P_1_* state in the 2PB labelling step. Under the stated labelling conditions, 2PB selectively labelled oxidised cysteines to near completion. To prevent it smearing bands by interacting with SDS, we validated the chemical removal of unreacted 2PB. These experiments demonstrated that the selective 2PB labelling condition, that is *P_1_*, is met before SDS-PAGE.

**Figure 4.**
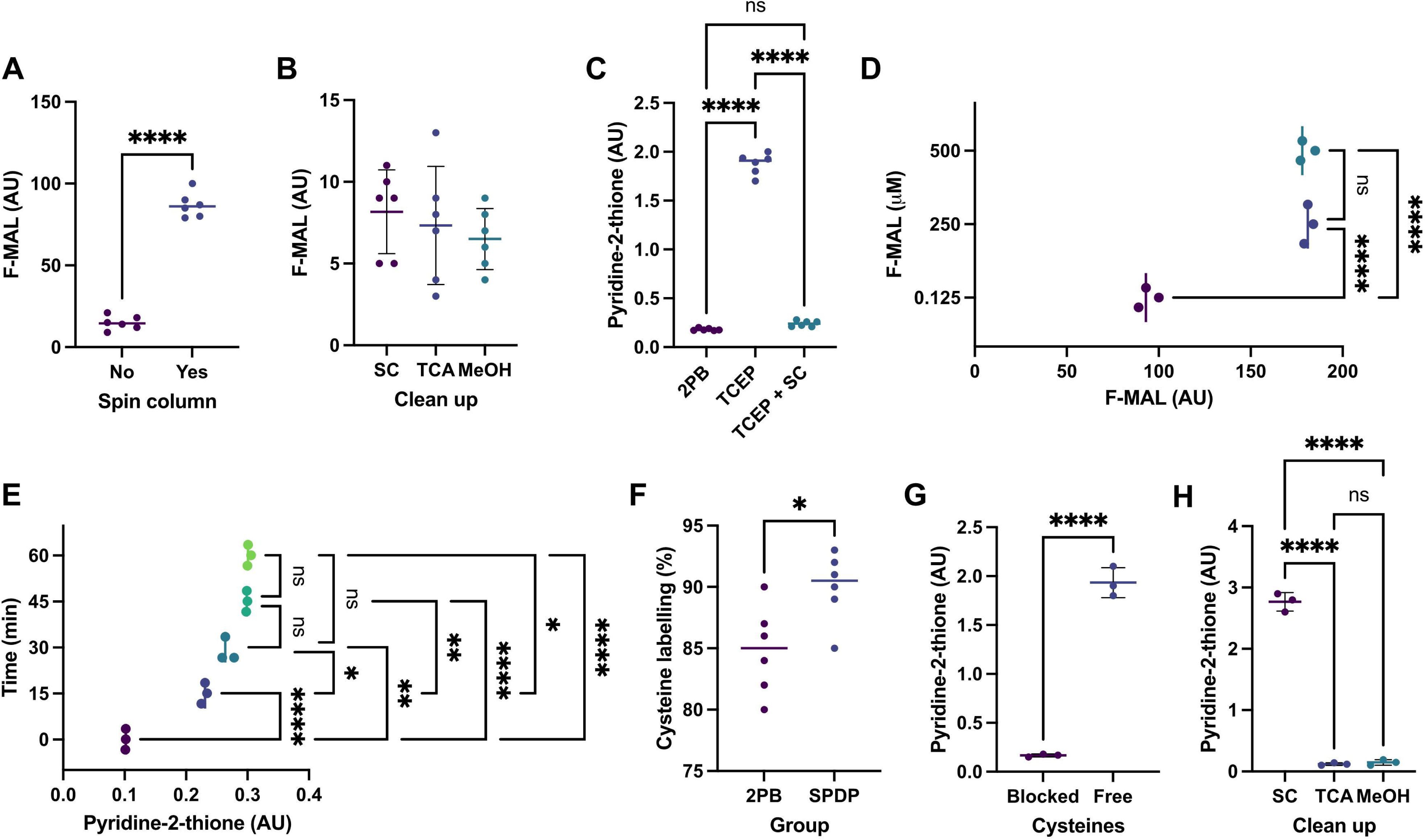
Validating the cysteine labelling procedures for 2PB-based Cleland immunoblotting. A. Significant difference in the fluorescent maleimide (F-MAL) signal (arbitrary units, AU) emanating from an *N*-ethylmaleimide (NEM)-alkylated tris-(2-carboxyethylphosphine) (TCEP)-reduced *X. laevis* lysates with (*n* = 6) compared to without (*n* = 6) a spin column (SC) step. This result indicates that the SC removes excess NEM from the sample. B. No significant differences in the F-MAL signal emanating from NEM-alkylated TCEP-reduced *X. laevis* lysates (*n* = 6 per group) cleaned up with a SC, trichloroacetic acid (TCA), or methanol (MeOH). This result validated that all the NEM is removed from the sample. C. No significant pyridine-2-thione release is observed from 2PB when TCEP is removed by an SC (*n* = 6 per group). This result validated our ability to remove TCEP form the sample, ruling out the possibility that unremoved reagent reduced newly formed disulfide bonds between 2PB and oxidised cysteine residues. D. No significant increase in cysteine labelling from 250 to 500 µM F-MAL in TCEP-reduced *X. laevis* lysates (*n* = 3 per group). To promote labelling, we added a molar excess of 2PB over the amount of cysteine in the sample, which is below 250 µM as estimated by an F-MAL titration curve. E. No significant increase in the extent of 2PB labelling, as determined via pyridine-2-thione release, in TCEP-reduced *X. laevis* lysates after 45 min (*n* = 3 per group). This time-course labelling experiment revealed that no appreciable pyridine-2-thione release occurred 45 min after adding 2PB to TCEP-reduced lysates. F. A significant difference in the extent of cysteine labelling, determined via a pulse chase experiment using an F-MAL standard curve (see methods), with SPDP (succinimidyl 3-(2-pyridyldithio)propionate) vs. 2PB in TCEP-reduced *X. laevis* (*n* = 6 per group). This experiment demonstrated the existence of a steric penalty imposed by the PEG-payload on the ability of 2PB to label cysteine residues. G. No appreciable pyridine-2-thione in NEM-blocked compared to TCEP-reduced *X. laevis* lysates (*n* = 3 per group). This experiment demonstrated that 2PB labelling cysteine residue-specific. H. A significant difference in pyridine-2-thione from 2PB after an SC clean up compared to TCA or MeOH as determined after adding TCEP to NEM-alkylated *X. laevis* eluents or pellets (*n* = 3 per group). This experiment demonstrated that it is necessary to remove excess 2PB with TCA or MeOH before SDS-PAGE. The data in panels A, F, and G were analysed by an independent t-test. The data in panels B-E and H were analysed with a 1-way ANOVA with Tukey post-hoc testing as appropriate. In all panels: ns denotes P > 0.05 whereas *, **, ***, and **** denote P < 0.05, 0.01, 0.001, and 0.0001, respectively.

After validating the cysteine labelling procedures, we used the biotin tail of 2PB as a reaction reporter for experimentally testing *P_1_*via non-reducing “oxidising” streptavidin immunoblotting. Oxidising streptavidin immunoblotting unequivocally evidenced the *P_1_* state across a range of maximally labelled-2PB protein inputs, confirming the existence and stability of the C-S-S-P disulfide bond. In this regard, C-S-S-P stability should be similar to the bond dissociation energy (60 kcal mol^-1^) of a typical disulfide bond. This experiment validated that the *P_1_* condition is met, as demonstrated by the C-S-S-P-dependent biotin positive bands (Figure 5, left panel).

**Figure 5.**
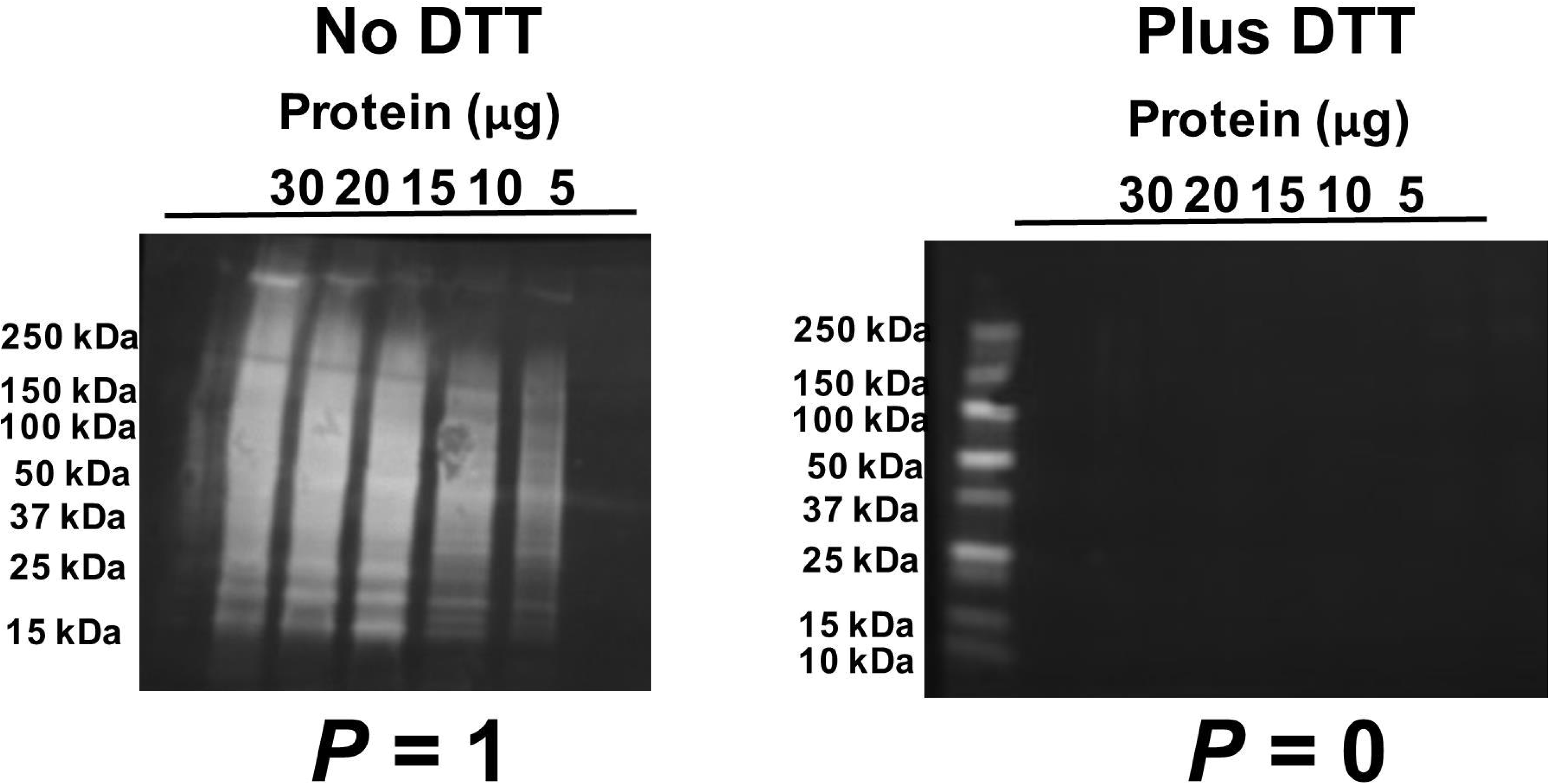
Experimental validation of the P-switch. Left. Oxidising (no DTT) streptavidin immunoblot image validating the presence of the PEG-payloads (*P* = 1) in proteins due to the presence of the biotin reporter in 2PB. In the *P_1_* state, the disulfide bond crosslinking 2PB to a cysteine residue is maintained. Right. Reducing (plus DTT) streptavidin immunoblot image validating the absence of the PEG-payloads (*P* = 0) in proteins due to DTT-mediated disulfide exchange reactions (see reactions 3-4). In the *P_0_* state, cysteines residues have been regenerated to the thiol/thiolate. These experiments were with fully 2PB-labelled *X. laevis* XI oocyte lysates.

To determine whether we could achieve the uniform *P_0_* state by reducing the C-S-S-P disulfide bonds, we soaked the gel in transfer buffer containing 324 mM DTT [20]. Per the Henderson-Hasselbalch equation, C-S-S-P reduction is favoured by the S^-^/SH ratio of the two sulfur atoms (p*K*_a_ = 9.2 and 10.1) in DTT being 1/8 and 1/63, respectively, at pH 8.3. In agreement with their Nernst equation predicted reducing power of -0.1332 and -0.3663 volts when [DTT] = 324 mM, reducing streptavidin immunoblotting confirmed the complete reduction of the C-S-S-P disulfide bonds per reactions 3 and 4. No biotin-positive bands were observed (Figure 5, right panel). This experiment validated that the essential uniform *P_0_*state condition is met.

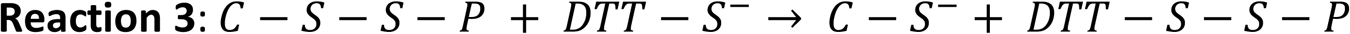

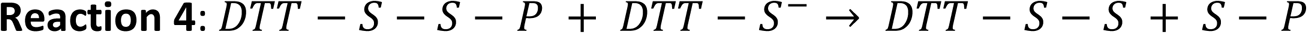

In sum, we experimentally validated that the ability to flick the P-switch between the electrical SDS-PAGE and transfer blot modules of the Cleland immunoblotting method.

### 2.4. The formal proposition is valid: All antigens were detected regardless of their position on the membrane under the uniform *P_0_* condition

After validating the P-switch, which is the foundational deductive argument upon which it rests, we experimentally tested the formal proposition that when *P_0_* = *A_1_* Cleland immunoblotting will invariably succeed. To do so, we measured <u>cdc20</u> in maximally 2PB-labelled *Xenopus laevis* (*X. laevis*) oocytes, for the reasons set out in section 2.5, in the *P_1_* and *P_0_*states. To detect cdc20 cysteine redox proteoforms, the *P_1_* state is needed as no endogenous mobility-shifted bands were observed in non-reducing SDS-PAGE (Figure 6A-B).

**Figure 6.**
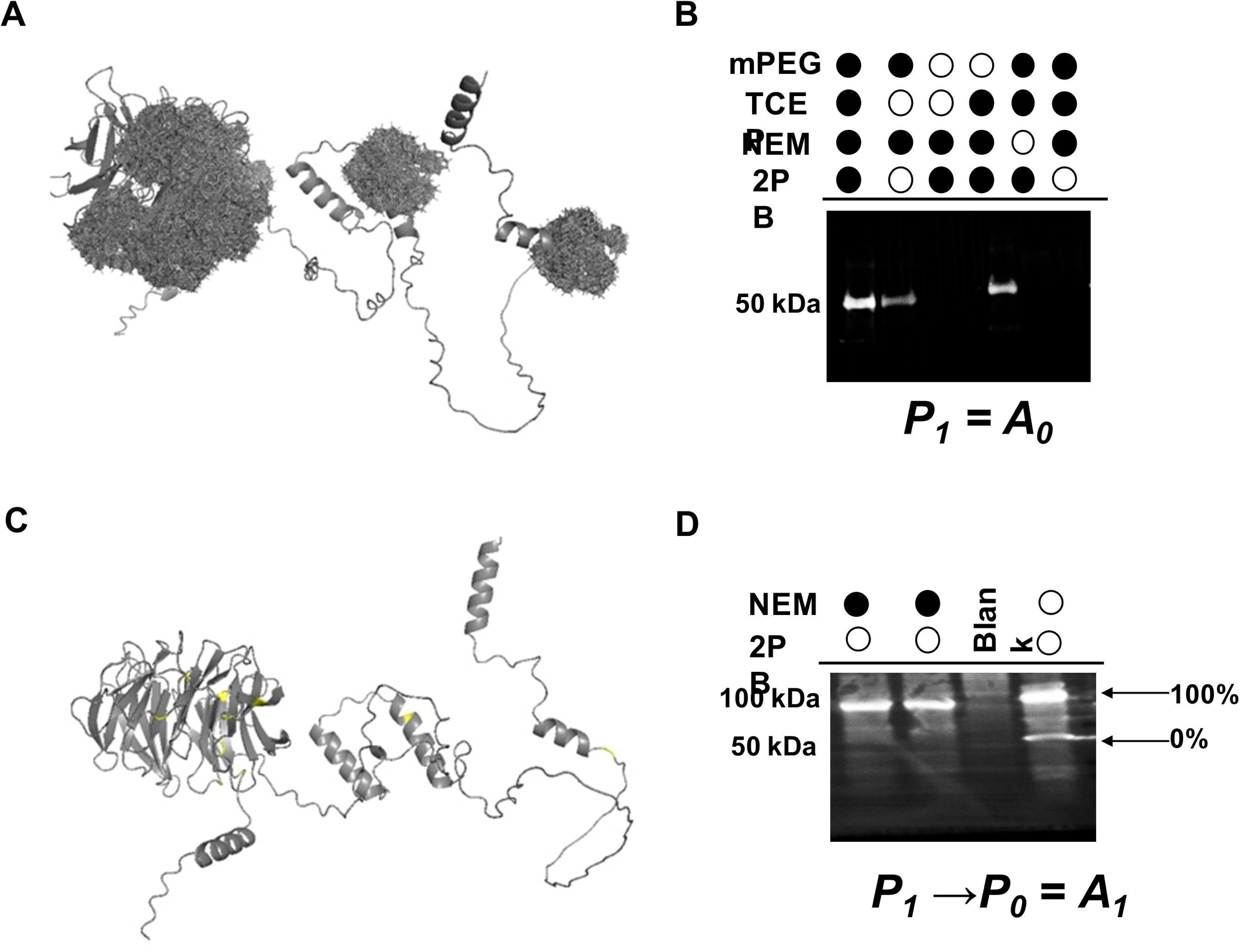
The formal proposition is valid: All antigens were detected regardless of their position on the membrane under the uniform *P_0_* condition. A. Image displaying cdc20 in the fully PEG-conjugated state (*P* = 1). B. Cdc20 immunoblot image under oxidising (no DTT) conditions where *P* = 1. White-filled circles denote the cysteine-reactive label. Lane 1 = no label, Lane 2 = 2PB labelled & TCEP reduced, Lane 3 = mPEG2 labelled, Lane 4 = mPEG2 labelled & TCEP reduced, lane 5 =, and lane 6 = 2PB labelled. The antibody failed to bind cdc20 in 2PB labelled lysates (lane 6) under oxidising conditions. Hence, *P_1_* = *A_0_*. C. Image showing cdc20 in the PEG-free state, with the free cysteine groups coloured yellow. D. Cdc20 Cleland immunoblot image. Lane 1-2 = ‘fully’ 2PB labelled, and Lane 3 = 5 µg (reduced) NEM-alkylated spiked into 15 µg 2PB (oxidised) labelled sample. Reducing the gel DTT enabled the antibody to bind cdc20. Hence, *P_1_* →*P_0_* = *A_1_*. These experiments were performed with maximally 2PB-labelled *X. laevis* lysates. Unless otherwise stated, the protein loading amount = 20 µg. An amount validated to remove the PEG-payloads (Figure 5).

Irrespective of whether we used the conventional m-PEG assay thioether bond or our novel C-S-S-P disulfide bond to conjugate iso-mass (2-3 kDa) PEG-payloads to cysteines residues, no cdc20-specific bands were detected under the *P_1_* condition. As visualised in Figure 6A, the PEG-payloads masked a substantial portion of cdc20, which would explain the lack of antibody binding. Indeed, we detected cdc20 in NEM alkylated lysates. NEM forms a thioether bond with the cysteine thiolate without introducing a PEG-payload. Tellingly, we detected cdc20 when the C-S-S-P disulfide bonds were reduced with TCEP before SDS-PAGE. Hence, *P_1_* = *A_0_*when cdc20 is the antigen.

We released the C-S-S-P-dependent PEG-payload from mobility-shifted cysteine redox proteoforms by soaking the gel in DTT to achieve the essential *P_0_* condition (Figure 6C). Under the *P_0_* condition, the antibody detected cdc20 regardless of the position of the antigen on the membrane. As displayed in Figure 6C, all of cdc20 is available for antibody binding in the *P* = 0 state. Since PEG-conjugates experience greater friction as they migrate through the porous gel matrix under an applied electromagnetic field [5], the position of the mobility-shifted band was ≈100 rather than ≈75 kDa. This result proved the formal proposition that for cdc20 *P_success_* is 1 when *P_0_*, irrespective of the position of the antigen (Figure 6D).

In sum, a novel cysteine redox sequence enabled Cleland immunoblotting to detect hitherto intractable cysteine redox proteoforms, proving the formal proposition for cdc20. The performance improvement was total: the m-PEG assay failed to detect cdc20 (0%-effective) whereas Cleland immunoblotting detected cdc20 (100%-effective).

### 2.5. Cleland immunoblotting unmasked unexpected cysteine redox proteoforms

After proving the formal proposition for cdc20, we used Cleland immunoblotting to study the redox dynamics of cdc20-specific cysteine redox proteoforms during fertilisation in *X. laevis* for several reasons. First, cdc20 plays a pivotal role in the oocyte-to-zygote transition by regulating meiosis and mitosis [23]. Second, cdc20 forms proteoforms [24], but cysteine redox proteoforms are unstudied. Third, we found that *X. laevis* oocytes house a vast biologically accessible theoretical *i* space, comprising up to 2.2 x 10^18^ unique cysteine redox proteoforms (Table 2, and supplementary figure 2).

**Table 2.**
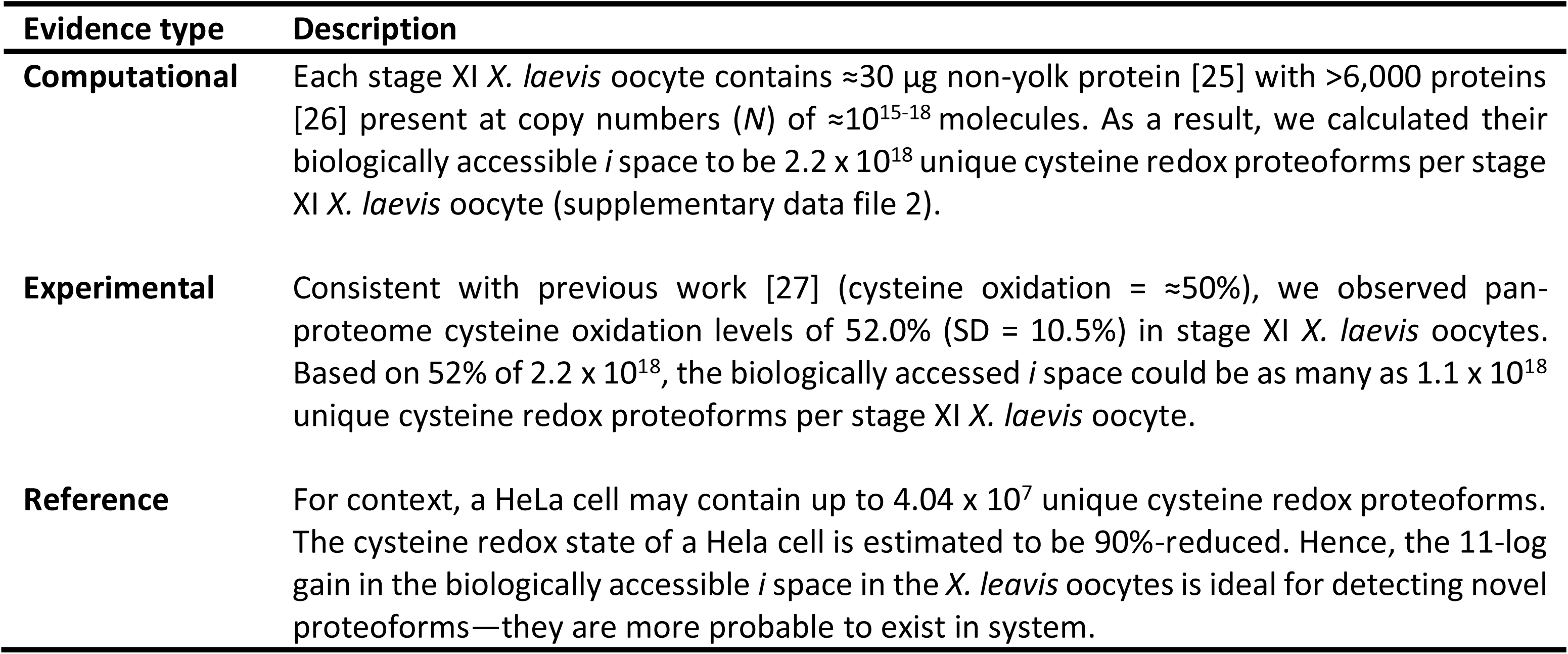
A brief description of the different lines of scientific reasoning, evidence types, that rationally informed the selection of the *X. laevis* oocytes model system. Supplementary Figure 2 and saupplementary data file 2 contains further information.

Mathematically, the 10 cysteine residues in cdc20—Cys-37, 105, 321, 346, 373, 397, 401, 412, 465, 481—can form 11 percentage redox graded bands (0-100%-oxidised). The geometric band proteoform structure for the 1,024 possible cdc20-specific cysteine redox proteoforms is as follows:

|  |  |  |  |  |  |  |  |  |  |  |  |
| --- | --- | --- | --- | --- | --- | --- | --- | --- | --- | --- | --- |
| % | 0 | 10 | 20 | 30 | 40 | 50 | 60 | 70 | 80 | 90 | 100 |
| *k* | 0 | 1 | 2 | 3 | 4 | 5 | 6 | 7 | 8 | 9 | 10 |
| *i* | 1 | 10 | 45 | 120 | 210 | 252 | 210 | 120 | 45 | 10 | 1 |

How many of these 1,024 unique cysteine redox proteoforms are biologically accessed? Unexpectedly, Cleland immunoblotting revealed that just 2 out of 1,024 (0.19%) unique cysteine redox proteoforms are biologically accessed before and after fertilisation in *X. laevis* (Figure 7). The unique, class 1, assigned coordinates mapped to the binary 100%-reduced and 100%-oxidised cysteine redox proteoforms, respectively. The mobility-shift displayed by the 100%-oxidised form was identical to the maximally 2PB-labelled form derived from the same samples. Hence, the cysteine redox state of cdc20 is a function of two binary proteoforms.

**Figure 7.**
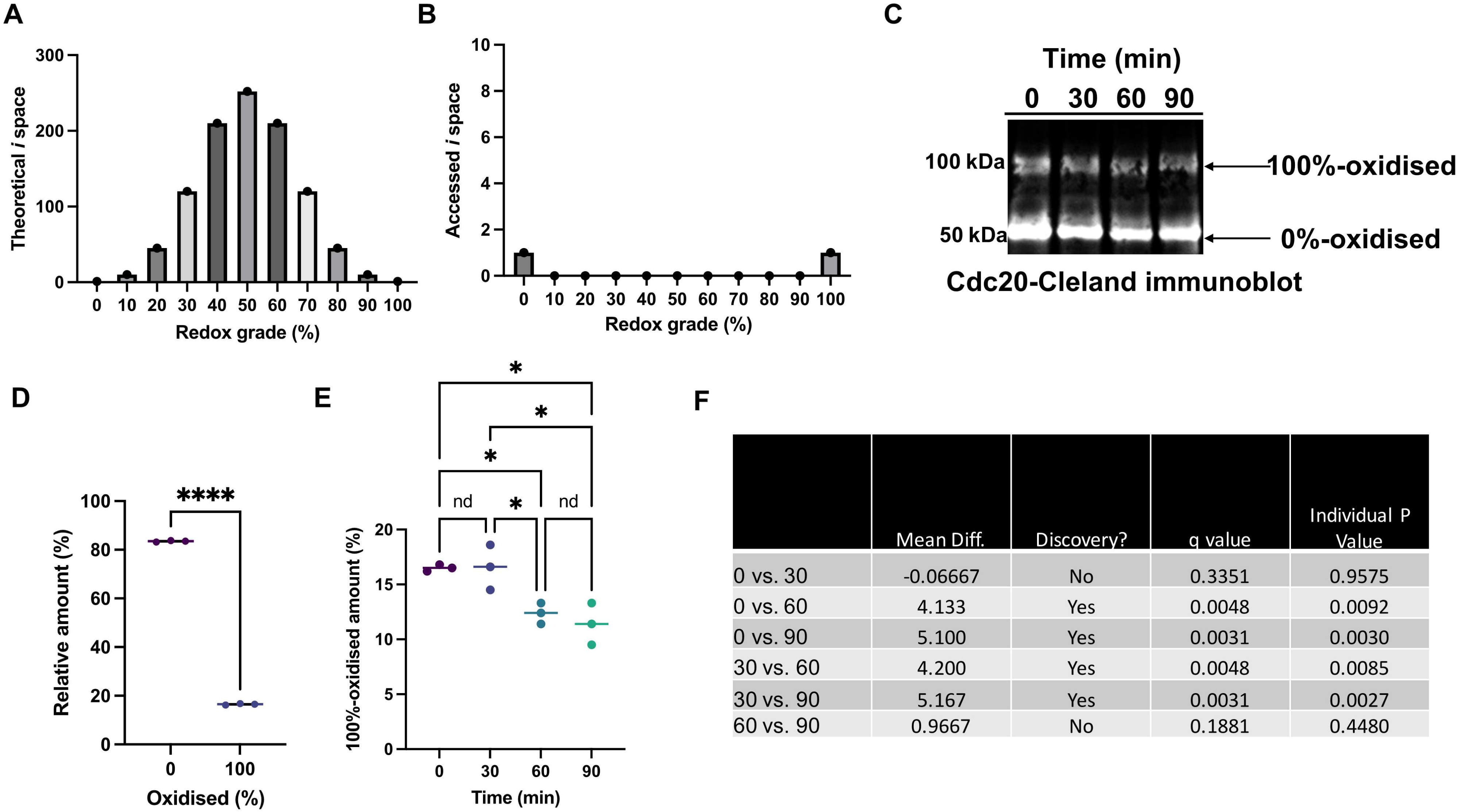
Cleland immunoblotting unmasked unexpected cdc20-specific cysteine redox proteoforms. A. Histogram of the number of theoretical cysteine redox proteoforms (*i* space) per percentage redox grade. The percentage redox grade being specified by the *k* value (e.g., 0% = 0 oxidised cysteines). B. Plot of the biologically accessed *i* space, showing that two (0.195%) out of 1,024 cysteine redox proteoforms were present. C. Immunoblot image showing a representative cdc20 Cleland immunoblot experiment (protein loading input = 15 µg per lane). The time indicates the sampling point where 0-min denotes the stage XI *X. laevis* oocyte and 30-, 60, 90-min denote the times that 1-cell zygotes were collected post-fertilisation. The text annotation denotes the positions of the 0 and 100%-oxidised cdc20-specific cysteine redox proteoforms. D. A plot displaying the relative amount of the 0 and 100%-oxidised cdc20-specific cysteine redox proteoforms in the stage XI *X. laevis* oocytes. A significant difference between the reduced (M = 83.5%, SD = 0.3%) and oxidised (M = 16.5%, SD = 0.3%) states was observed according to a paired t-test (*t*(2) = 0.193.4, *p* = <0.0001, mean difference = 67.0%). E. A plot displaying the relative amount of the 100%-oxidised cdc20 proteoform as determined by Cleland immunoblotting at each sampling time point (*n* = 3). F. A table mapping the statistical significance annotations in plot E to discoveries (yes or no) as determined by a one-way ANOVA (*F*(3,8) = 10.8, *p* = 0.0044). with post-hoc testing and corrections for multiple comparisons. Mean differences, q values and individual *P* values are shown.

Internally normalised analysis of their relative abundance revealed that the 100%-oxidised cdc20-proteoform comprised 16.5% (SD = 0.3%) of the total signal in oocytes (0-min). Interestingly, the relative amount of the 100%-oxidised proteoform remained constant between 0 and 30-min post-fertilisation (mean difference = 0.07%). However, the amount of the 100%-oxidised proteoform decreased at 60-min (-4.1%) and 90-min (-5.1%) post-fertilisation vs. 0-min. For example, it decreased from 16.5% (SD = 0.3%) at 0-min to 11.4% (SD = 1.4%) at 90-min post-fertilisation (supplementary Figure 3). The redox dynamics of cdc20 toward a more reduced state were attributable to a decrease in the relative amount of the 100%-oxidised proteoform.

More generally, *X. laevis* oocytes are ripe for harvesting novel data: Many more display cysteine redox proteoform dynamics because fertilisation-decreased pan-proteome cysteine oxidation levels by approximately 10% (supplementary Figure 4). In sum, Cleland immunoblotting derived novel insight by unmasking a redox-sensitive protein: the cysteine redox state of cdc20, as encoded by binary proteoforms, responded to fertilisation.

### 2.7. Qualifying the extent of the advance

Could a 100%-oxidised cdc20-specific proteoform be discovered another way? Unlikely. Per section 2.4, the m-PEG assay cannot detect cdc20. What about alternate immunological methods? No. While RedoxiFluor [19] quantified cdc20 cysteine redox state and recapitulated the fertilisation-induced decreased in oxidation (Figure 8), it cannot measure specific proteoforms. Bar monothiols, there is no known way to reconstruct the proteoform unit from the oxidised peptide parts in BU-MS. BU-MS illustrated the novelty of our results. Dividing the relative amount of the 100%-oxidised proteoform by *r* (i.e., 16.5%/10) would yield an oxidation value of 1.65% per cysteine residue. A value of 1.65% oxidised would not be classed as a discovery in BU-MS using any current interpretational rubric. The proteoform basis can unmask discoveries that would otherwise be overlooked at the peptide basis.

**Figure 8.**
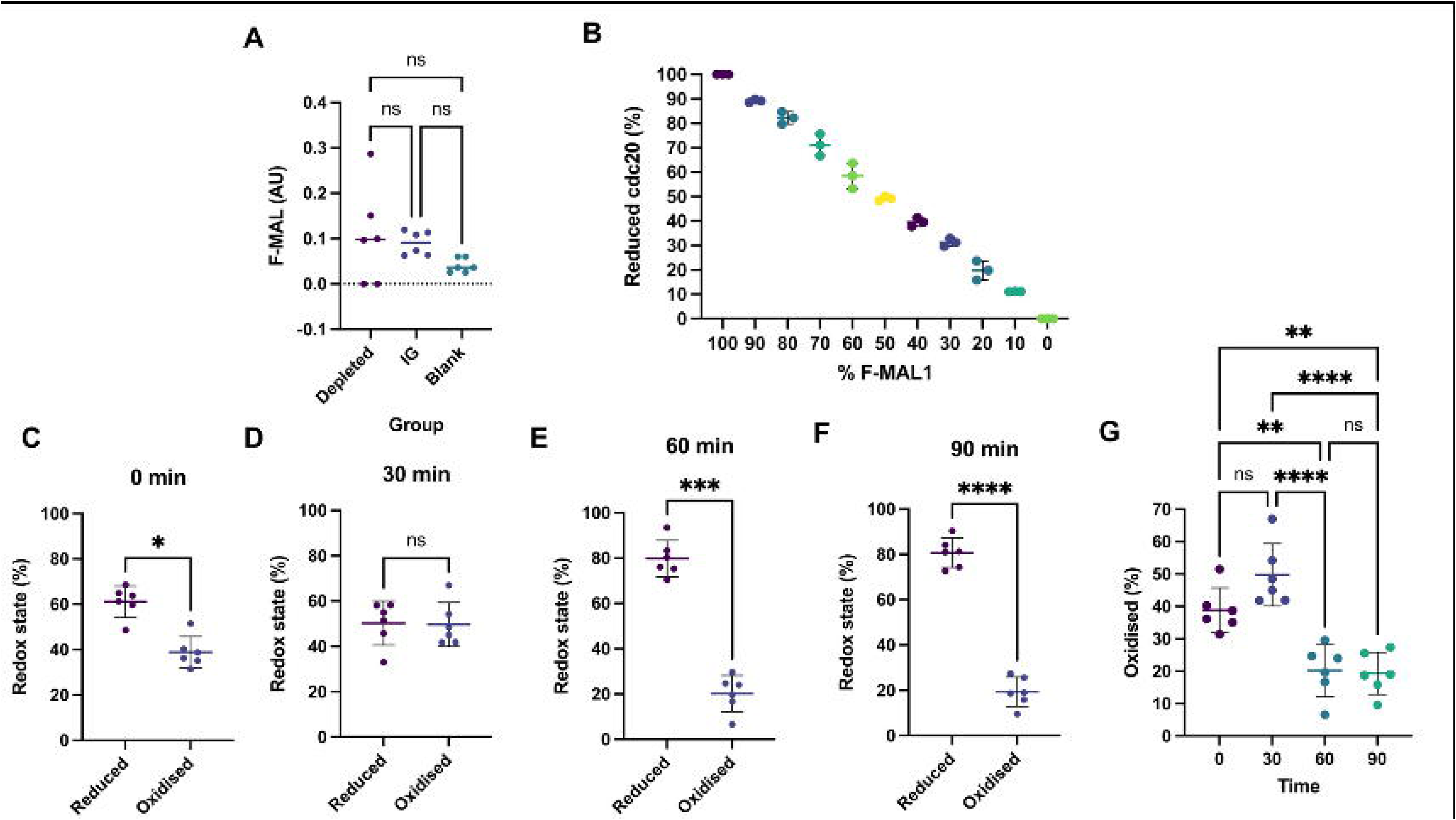
Protein A mode cdc20 RedoxiFluor assay. A. The cdc20 antibody specificity test displaying the fluorescent maleimide (F-MAL) value in arbitrary units (AU) for an immunoglobin control, cdc20-(immuno)depleted, and assay blanks. There were no significant differences between any of the conditions as determined by a one-way ANOVA. This result indicates that the assay signal derives from cdc20 itself and/or any naturally interacting proteins. B. An example assay calibration curve as prepared using standards. C-F. Plots displaying the cysteine redox state of cdc20 before (0-min) and at various time-points after fertilisation as determined using protein A mode RedoxiFluor. Statistical significances were assessed using paired *t*-tests. G. A plot displaying the percentage cysteine oxidation of cdc20 over time. Statistical significance was assessed using a one-way ANOVA with Tukey post-hoc testing. In all panels: ns denotes P > 0.05 whereas *, **, ***, and **** denote P < 0.05, 0.01, 0.001, and 0.0001, respectively.

Based on the following probability (*P*) values:

- *P* of ionising cdc20 = 0.6
- *P* of selecting cdc20 for MS = 0.5
- *P* of fragmenting cdc20 = 0.8
- *P* of detecting all cysteine peptides = 0.7

And without considering spectral interference from other proteoforms [28], the probability of TD-MS making the discovery is 16.8% (*P* = 0.6 x 0.5 x 0.8 x 0.7 = 0.168). While the chance is non-zero, it is unlikely that TD-MS would have discovered a 100%-oxidised cdc20-specific proteoform. Still, the chances improve for class 2 assignments. Of interest, the “isobaric” residues and the “isobaric” clusters of proteoforms in bands are ostensibly different manifestations of the same thing. They underpin 2A and 2B classifications in TD-MS and Cleland immunoblotting, respectively [3]. Irrespective of the method-dependent *P* values for a given proteoform assignment, one would only need standard lab equipment to make the discovery using Cleland immunoblotting.

In sum, we would have had between 0 and 16.8% chance of discovering a 100%-oxidised cdc20-specific proteoform using other methods. The extent of the advance made by Cleland immunoblotting, especially for those without access to an ion cyclotron resonance instrument for TD-MS [29], is considerable.

### 2.7. On the novel insights derived by Cleland immunoblotting

By identifying a fertilisation-responsive protein, Cleland immunoblotting derived novel insight into the cysteine redox state of cdc20. This result paves the way to determine whether cdc20 is redox regulated. As Cleland immunoblotting demonstrated, the cysteine redox state of the cdc20 molecules (R_cdc20_) is a function of 2 binary proteoforms per equation 8. To illustrate the non-obvious nature of this novel finding, the <u>Solution_Space_Cal.py</u> script estimated that there could be 3.03 x 10^16^ possible solutions to equation 3 for cdc20 for a population of just 196 molecules (supplementary data file 3). The estimated size of the measured cdc20 population is in the order of 10^16^ molecules.

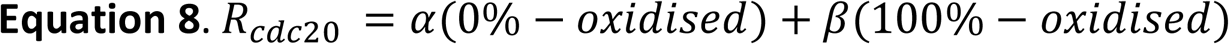

Equation 8 demonstrated that the percentage redox state of a population of protein molecules is governed by specific cysteine redox proteoforms. By extension, the cysteine redox state of the proteome is a function of proteoforms, which underscored how cysteine redox biology is rooted in the dynamics of specific proteoforms. Indeed, every protein with a cysteine is a cysteine redox proteoform (equation 2). Per equation 8, R_cdc20_ is not simply the result of any one oxidised cysteine residue in isolation. Rather, it is the product of oxidised residues in the context of a specific proteoforms—100%-reduced and 100%-oxidised molecules.

From a statistical mechanics perspective, the discovery of a 100%-oxidised cdc20 proteoform was upending. The bimodal, symmetrical distribution of cdc20 into 2 diametrically opposite cysteine redox proteoform states was thermodynamically improbable. The probability of ≈2.9 x 10^16^ cdc20 molecules being either fully reduced or fully oxidised is significantly lower than being in a partially oxidised state. There are 1,022 ways for cdc20 molecules to be partially oxidised vs. only 2 ways to be either fully reduced or fully oxidised, resulting in a probability ratio of ≈3-logs (1,022:2). Equation 9 revealed that the odds of observing a binary state are 1:500 (*P* = 0.002, or 0.2%).

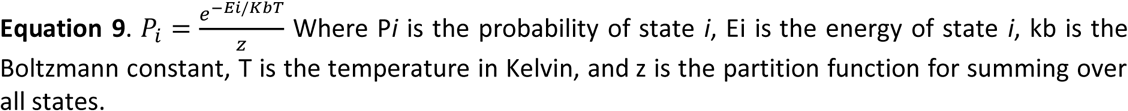

Unmasking a 100%-oxidised proteoform was unexpected as no known structural or kinetic arguments favour cdc20 oxidation. For example, cdc20 lacks structural disulfide bonds (Figure 6A&C). Moreover, the rate (*k*) constant with hydrogen peroxide (H_2_O_2_) to yield a sulfenic acid (RSOH) is likely to be similar to most cysteines: *k* = 1-10 M^-1^ s^-1^ at 37°C [30]. Based on the following parameters:

- Cdc20 *N* = 2.90 x 10^16^ in an *X. laevis* oocyte [26]
- cdc20 cysteines = 2.90 x 10^17^ (i.e., *N*r*)
- *N* 100%-oxidised cdc20 = 4.35 x 10^15^ (15%)
- *N* 100%-oxidised cysteines = 4.35 x 10^16^
- [H_2_O_2_] = 10 nM
- *k* = 5 M^-1^ s^-1^ at 22°C.

It would take 150.2 billion years to oxidise 15% of the cdc20 molecules to the 100%-oxidised proteoform, far exceeding the estimated age of the universe. Hence, the formation of the 100%-oxidised form proceeds via a concerted, deliberate mechanism. The mechanism is likely unconventional. For example, the indiscriminate chemical reactivity of the hydroxyl radical could be selectively channelled to cysteines via a transition metal ion conduit. The conduit would react with H_2_O_2_ to produce the hydroxyl radical, directing it toward proximal cysteine residues. With a *k* of 100 or 1,000 M^-1^ s^-1^ for the “Fenton” type reaction [31], the necessary cysteine oxidation could be achieved in 4.1 to 41.1 hours, respectively. Reactions 5-8 describe one possible mechanism where cysteine-glutathione adducts are formed at the end [32]. The formation of superoxide in the process may amplify cysteine oxidation [33]. Plausible alternatives include phase separation and enzyme catalysis [34,35]. It is fitting that the unmasking of an unexpected proteoform requires, by any reasonable back of the envelope calculations, unconventional cysteine oxidation mechanisms.

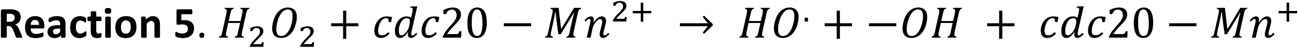

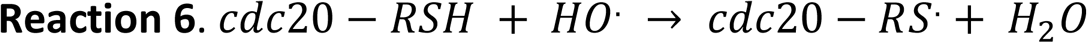

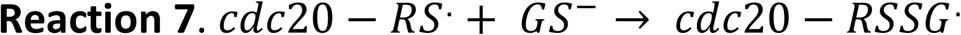

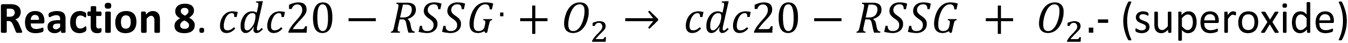

A fascinating feature of these mechanisms is the absence of intermediary partially reduced/oxidised states. Partially reduced/oxidised forms must be formed along the processive 10-reaction redox pathway between the binary states without remaining there long enough to be detected (*Ni*-10-90% *t* = <30-min). One or more factors must act on them to quickly complete the 10-reaction sequence and/or prevent any intermediates accumulating to a detectable level. The fact that a 10-reaction sequence proceeded to completion without intermediates defined a strong argument against adventitious oxidation during the sample work up. It would be fascinating to know which of the 1,022 possible routes between the binary proteoforms are traversed (Figure 9). Mapping them could decipher proteoform-encoded rules, such as cysteine oxidation dependencies [2].

**Figure 9.**
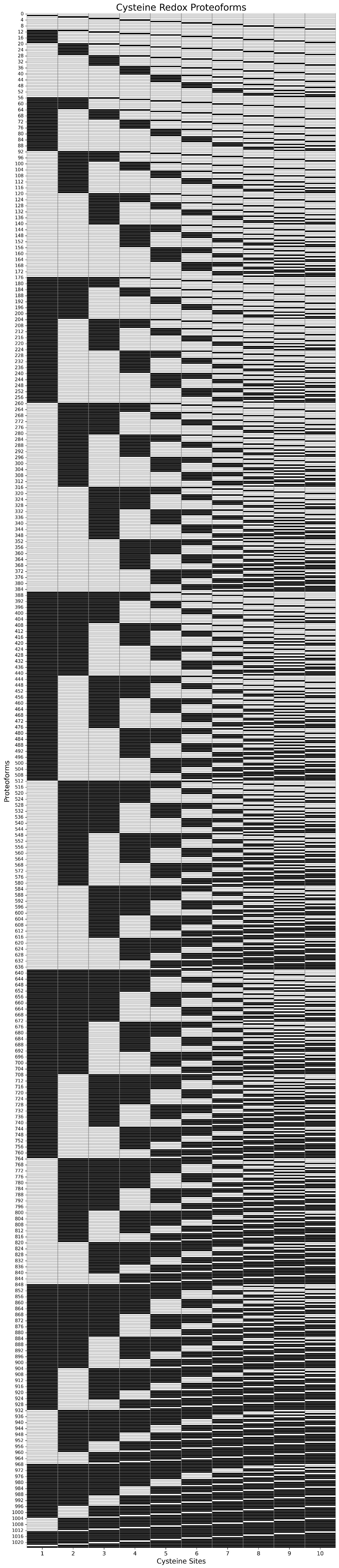
A plot of illustrating 1,024 theoretical cysteine redox proteoforms for a protein with 10 cysteine residues, such as cdc20. Each column corresponds to a residue and each row maps to a specific proteoform. The redox state is encoded in binary where grey and black denote the reduced and oxidised redox states, respectively. The proteoforms are ordered by their *k* value. For example, *k*_0 = 0%-oxidised (0,0,0,0,0,0,0,0,0,0). The plot was generated using the <u>proteoform_map.py</u> script.

A minimum of 9 other proteoforms, one for each grade, must be formed to explain our results. At least 1.1% of the abstract cdc20-specific *i* space is accessed. The principle of least action suggests that out of the 9 to 1,022 possible pathways to traverse the percent redox grades, the system adopts the pathway that minimises the overall energy and time required to do so. This is consistent with coordinated and rapid redox reactions, facilitated by specific catalysts or reaction environments that ensure intermediates are short-lived. On energy, cysteine redox proteoforms can be related to entropy (*S*) via equation 10. The kinetically feasible transitions between thermodynamically improbable high-energy states, such as a 100%-oxidised proteoform, suggest a way to satisfy *S* by increasing disorder using redox couples over the long lifespan of the oocyte.

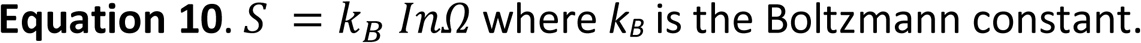

In sum, novel Cleland immunoblotting-derived insights opened new avenues of biological discovery.

### 2.8. The Cleland immunoblotting conjecture

Irrespective of the position of the antigen on the membrane, the formal proposition that *P_success_* is 1 when *P* = 0 was proved for cdc20. Beyond cdc20, we conjecture that the same deductive principles underpinning Cleland immunoblotting will logically guarantee a successful outcome for other proteins, provided that essential conditions, such as achieving the uniform *P_0_* state, are met. In this regard, vast tracts of the human proteome have been measured by immunoblotting, meaning *A_1_* is already evidenced for most proteins when *P* = 0.

We envisage that Cleland immunoblotting will advance cysteine redox proteoform research. For example, by quantifying how many of the 281,482 possible percentage redox graded bands theoretically available to 19,817 proteins and their 262,025 cysteine residues are biologically accessed in humans (supplementary data file 4). To support such endeavours, we defined the molecular mass (kDa) of each <u>human reference proteome</u> FASTA file protein entry in the 100%-reduced and 100%-oxidised (100%-oxidised = 100%-reduced + [*r*\*5 kDa]) *i*-states using the <u>Cleland_Cys_Proteome.py</u> script (supplementary data file 5). We estimate that approximately 70 and 80% of the 19,817 proteins in the human cysteine proteome would be detected by Cleland immunoblotting with a gel resolution range of 150 and 200 kDa, respectively (Figure 10). For larger proteins with high cysteine residue integers, 2-dimensional versions (<u>2D-Cleland immunoblotting</u>) can be used. As a technology that should be capable of detecting the vast majority of proteins in the human cysteine proteome, Cleland immunoblotting may become a “cheat code” for measuring oxidative stress [36].

**Figure 10.**
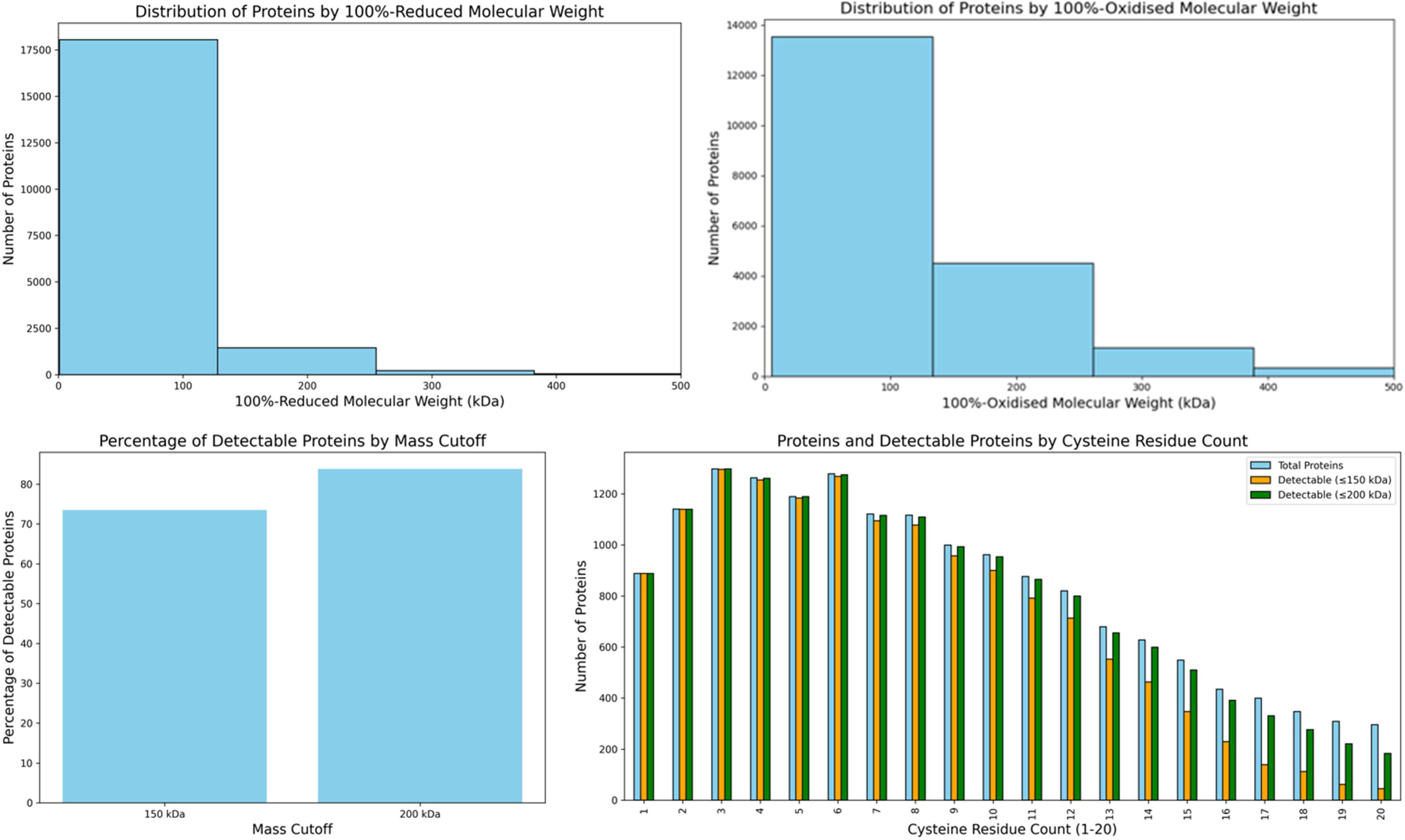
Estimating the number of human proteins that Cleland immunoblotting can detect. Top. Histograms distribution of the cysteine human proteome (19,817 proteins) by molecular weight in the 100% reduced (left) and 100%-oxidised (right) *i*-states. Bottom. Left. Plot showing the percentage of proteins estimated to be detectable by Cleland immunoblotting with a 150 and 200 kDa cut-off. Right. Total proteins and the number of detectable proteins in the 150 and 200 kDa cut-off range for cysteine residue integer values 1-20. Plots were generated using the <u>Cys_Proteome_Analysis.py</u> script.

To provide a resource for implementing Cleland immunoblotting, the clelandimmunoblotting.streamlit.app/ can visualise the band profiles of proteins of interest using their UniProt accessions (Figure 11).

**Figure 11.**
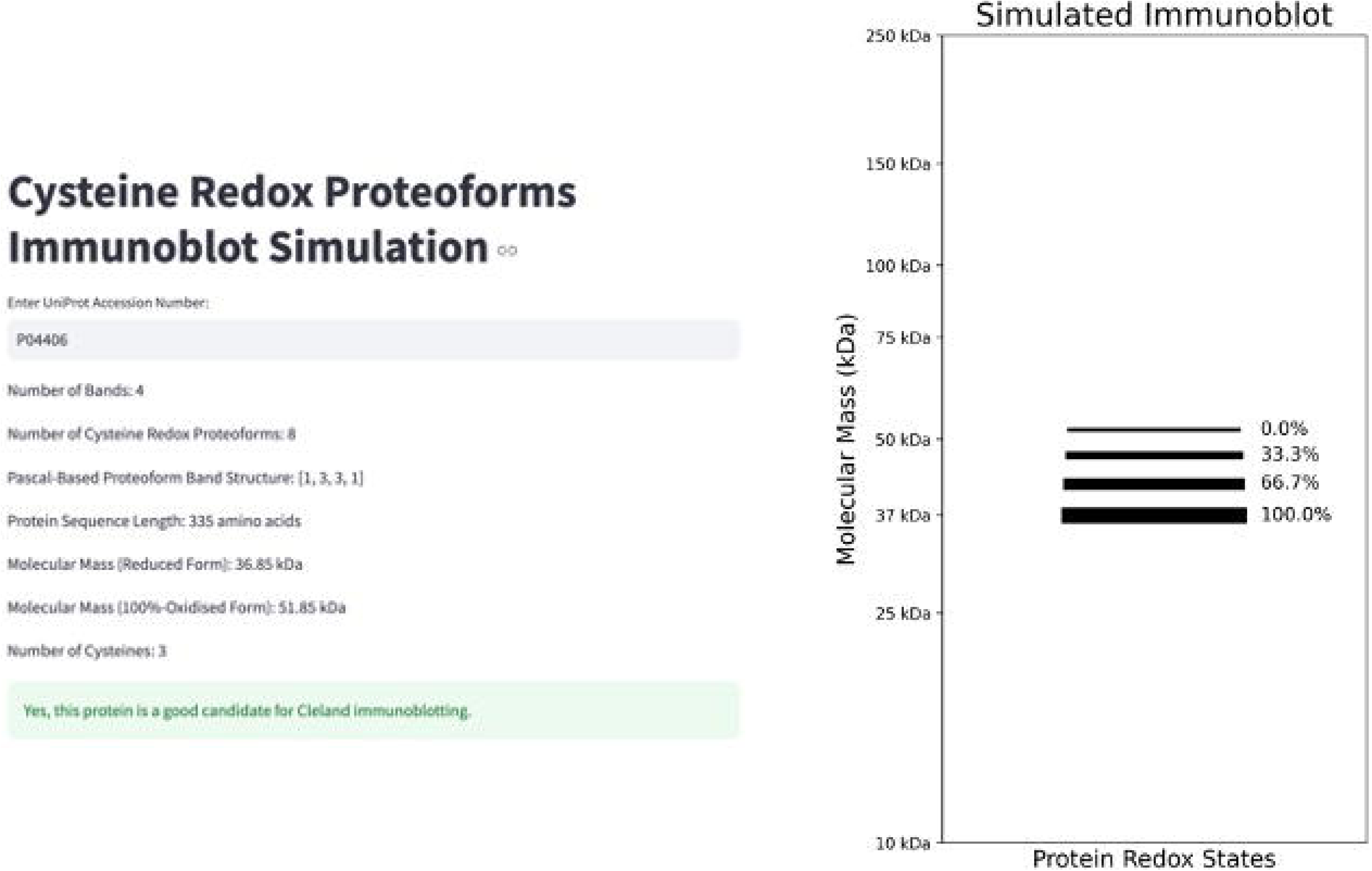
Cleland immunoblotting streamlit application. Left. A screenshot print output for a UniProt accession (GAPDH = P04406) from the <u>clelandimmunoblotting.streamlit.app/</u>, which was created using the <u>Cleland_app.py</u> script. The automated print output lists the number of bands (equation 5), the number of cysteine redox proteoforms (equation 1), and the Pascal-based geometric bands structure (equation 7), number of cysteines, and the molecular mass of the reduced and 100%-oxidised proteoforms. Additionally, a message that indicates the suitability of the inputted candidate for Cleland immunoblotting is displayed. Right. The application simulates an immunoblot displaying the percentage redox graded bands (listed from 100% to 0% reduced for the protein of interest). The image can be downloaded from the app.

In sum, we conjecture that Cleland immunoblotting can detect cysteine redox proteoforms belonging to diverse proteins, provided the essential conditions are met.

## 3. Conclusion

We succeeded in overcoming longstanding technical challenges by developing Cleland immunoblotting technology for measuring cysteine redox proteoforms. In so doing, we derived novel findings and insights (Table 3). By delivering the technological means to map the uncharted cysteine redox proteoform landscape, we envisage that Cleland immunoblotting will open new avenues of discovery.

**Table 3.** Section and **type stratified summary of the main observations, findings, and/or insights of the present work.**

| Section | Type | Observation, finding, and/or insight |
| --- | --- | --- |
| 2.1 | *Theoretical* | The deductive argument that $P_1 \rightarrow P_0 = A_1$ defined the formal proposition that underpinned the development of the Cleland immunoblotting method. |
| 2.2 | *Resource* | Elaborated a novel 2PB reagent for realising the P-switch $P_1 \rightarrow P_0$. |
|  | *Theoretical* | Elaborated a chemistry-enabled method for testing the formal proposition called Cleland immunoblotting. |
|  | *Theoretical* | Presented the mathematical basis of the technique, including the Pascal proteoform structure of the bands, and a cysteine redox proteoform classification system. |
| 2.3 | *Technical* | Procedures for labelling oxidised cysteine residues with 2PB were validated. |
| | *Technical* | The P-switch was validated by oxidising ($P = 1$) and reducing ($P = 0$) streptavidin immunoblotting. |
| 2.4 | *Technical* | In the $P = 1$ state, cdc20 cannot be detected by immunoblotting. |
| | *Technical* | The formal proposition that $P_1 \rightarrow P_0 = A_1$ was validated for cdc20. |
| 2.5 | *Biological* | There could be up to $1.1 \times 10^{18}$ cysteine redox proteoforms with at least one oxidised cysteine residue in stage XI *X. laevis* oocytes. |
|  | *Biological* | Just 2 out of 1,024 cdc20-specific cysteine redox proteoforms are biologically accessed before and after fertilisation in *X. laevis*. |
|  | *Biological* | The accessed cysteine redox proteoforms mapped to the fully, all 10 cysteine residues, reduced and oxidised forms. |
|  | *Biological* | The relative amount of the 100%-oxidised proteoform decreased after fertilisation. |
| 2.6 | *Technical* | Other immunological techniques could not measure cdc20-specific cysteine redox proteoforms, limiting their ability to derive the insights described in section 2.5. |
|  | *Technical* | It is unlikely the insights described in section 2.5. could have been derived using top-down mass spectrometry. |
| 2.7 | *Theoretical* | From a statistical mechanics perspective, the bimodal, symmetrical, distribution of cdc20 into binary, fully reduced and oxidised, proteoforms was unlikely. |
|  | *Theoretical* | Kinetic arguments suggested that generating the 100%-oxidised proteoform likely requires unconventional cysteine oxidation mechanisms. |
| 2.8 | *Conjecture* | When the essential conditions are met, Cleland immunoblotting is conjectured to capable of detecting most proteins. |
|  | *Theoretical* | Cleland immunoblotting is estimated to be able to detect over 70% of the human cysteine redox proteome. |
|  | *Resource* | A streamlit Cleland immunoblotting application is available for automatically solving the relevant equations and generating simulated immunoblot outputs. |

## 4. Methods

### 4.1. Ethical approval and sample collection

After receiving institutional ethical approval (#ETH2021-0222), stage XI oocytes were collected from 2 adult *X. laevis* females housed in the <u>European *Xenopus* Resource Centre</u> at 18°C. The stage XI oocytes were fertilised per previous work [6], and the 1-cell zygotes were collected 30-, 60-, and 90-min post-fertilisation. Groups of 20 oocytes or 1-cell zygotes were collected per sample [37].

### 4.2. The redox state of the cysteine proteome

To measure the cysteine redox state of the proteome, we used global mode RedoxiFluor[19,38,39]. Reduced and reversibly oxidised cysteine residues were labelled with spectrally distinct fluorescin-5-maleimide (F-MAL1, ThermoFisher, UK, #62245) and AlexaFluor®647-C_2_-Maleimide (F-MAL2, ThermoFisher, UK, #A20347) fluorophores, respectively. The F-MAL1/2 labelled samples were measured in triplicate in a plate reader for 100 ms. The percentage cysteine redox state of each sample was calibrated using 0-100%-oxidised standards as previously described.

### 4.3. RedoxiFluor

To measured cdc20 cysteine oxidation, we used protein A mode RedoxiFluor, exploiting the proven ability of the selected monoclonal antibody (see below) to immunocapture *X. laevis* cdc20 [22]. In brief, 0.1 μg of antibody in binding buffer (50%: 0.05% Tween-20 in phosphate buffered saline [PBST]; 50% Superblock [ThermoFisher, UK, #37580]) was added to each well of a black protein A derivatised microplate (ThermoFisher, UK, #15155) for 1 h at room temperature (RT) at 350 rpm. The protein A plates can bind the mouse IG subtype used. Unbound antibody was washed away (3 × 2 min PBST washes at 400 rpm), before assay calibrants (i.e., 10–90% reversibly oxidised standards), controls (i.e., immunodepleted, see below), or samples were added in duplicate and incubated in the dark for 2 h at RT. After removing unbound sample, wells were washed (3 × 2 min PBST washes at 400rpm), rinsed in PBS to remove excess Tween-20, and incubated with 4% SDS for 15 min at RT [39]. After subtracting the background, cdc20 cysteine oxidation was calculated in percentages (e.g., oxidised = F-MAL2/[F-MAL1 +F-MAL2]*100) after correcting (c) the F-MAL1 and F-MAL2 signals for their different quantum yields (q) and extinction co-efficient (e) integers per: c-F-MAL = [(F-MAL∗q)/e].

### 4.4. 2PB synthesis

To synthesise 2PB in a simple 1-pot chemical reaction, SPDP-dPEG®_36_-NHS ester (Sigma, UK, #QBD10867) and NH_2_-PEG_11_-Biotin (ThermoFisher, UK, #26136) were mixed in a 1:3 ratio for at least 1 h at 24°C in 1X PBS (pH 7.2). NH_2_-PEG_11_-Biotin molecules were in excess to compete with the aqueous hydrolysis of NHS to an amine unreactive form. The reaction vials were mixed every min for the first 15 min and 5 min thereafter. The resultant 2PB aliquots were stored at -20°C.

### 4.5. Validating the cysteine labelling procedures

*Fig4A*. To determine whether the spin column removed excess NEM (ThermoFisher, UK, #23030) from *X. laevis* oocyte lysates, we performed a fluorescent cysteine chase experiment. To do so, we reduced the NEM-alkylated samples with 5 mM neutral-TCEP (Sigma, UK, #580561) for 30 min at RT before adding 1 mM fluorescent-maleimide (F-MAL, Fluorescin-5-maleimide, ThermoFisher, UK, #62245) at RT with or without a prior spin column (Bio-Rad, UK, #7326222) step. The resultant F-MAL signals were measured at 494 (excitation) and 518 (emission) nm for 100 ms on a plate reader. The raw fluorescent data were background subtracted.

*Fig4B*. To compare clean up methods, the F-MAL experiments described above were repeated after cleaning a NEM-alkylated sample up with a spin column, trichloroacetic acid (TCA) (Sigma, UK, #T6399), or methanol (MeOH, VWR, UK, #179957). For the TCA condition, TCA was added at a 1:4 to the samples. The samples were incubated for 10 min at 4°C before being centrifuged at 14,000 *g* for 5 min at RT. After removing the supernatant, the protein pellets were washed with ice-cold acetone (Sigma, UK, #48358), re-spun at 14,000 *g* for 5 min at RT, and resuspended in F-MAL supplemented lysis buffer. For the organic solvent condition, the proteins were precipitated by sequentially adding MeOH(4 parts)-chloroform(1 part, Sigma, UK, #650498)-ddH_2_O (3 parts) to the samples. The samples were centrifuged at 14,000 *g* for 1 min at RT. After removing the aqueous layer, MeOH was added to the protein flakes before they were centrifuged at 20,000 *g* for 5 min at RT. MeOH aspirated pellets were air-dried before being solubilised in F-MAL supplemented lysis buffer. For each clean up condition, 1 mM F-MAL was added for 30 min at RT and removed via a spin column step, which we previously validated to remove all of the F-MAL [19]. Background subtracted fluorescent signals were quantified per Figure 2A.

*Fig-4C*. To determine whether the spin column removed excess 5 mM TCEP, we monitored pyridine-2-thione release at 343 nm for 100 ms in a plate reader after adding 1 mM hydrolysed 2-pyridyldithiol-PEG_36_-NHS to the eluents. To avoid any confounding impact of the NHS group, the 2-pyridyldithiol-PEG_36_-NHS molecules were hydrolysed for 4 h at 37°C in PBS. As a positive control, TCEP was added to hydrolysed 2-pyridyldithiol-PEG_36_-NHS. Background pyridine-2-thione release from hydrolysed 2-pyridyldithiol-PEG_36_-NHS in the absence of cysteine and TCEP provided the negative control.

*Fig-4D*. To determine whether the added 2PB would be in excess of the labelable cysteine residues in *X. laevis* egg lysates, we performed an F-MAL titration curve. To do so, increasing amounts of F-MAL1 was added to 50 µg of 5 mM neutral TCEP-reduced sample until no significant increase in F-MAL labelling was observed. The excess F-MAL was removed via a spin column step, before the fluorescent signals from the sample aliquots were measured in a 96-well microplate on plate reader[37,38].

*Fig-4E*. To optimise the labelling time, 5 mM 2PB was added to 75 µg protein aliquots of ‘fully’ (i.e., NEM-free) TCEP-reduced *X. laevis* egg lysates for the stated times in the dark at 4°C. Cysteine labelling diagnostic pyridine-2-thione release was serially monitored at 343 nm on a plate reader for 100 ms in samples incubated in 96-well microplate.

*Fig-4F*. To perform the pulse chase cysteine labelling experiment, 5 mM 2PB was added to 75 µg protein aliquots of NEM-free TCEP-reduced *X. laevis* egg lysates for 1 h at 4°C. The excess 2PB was removed via TCA precipitation before the samples were chased with 1 mM F-MAL for 30 min at 4°C. As a PEG-payload size control, the degree of labelling was compared to hydrolysed succinimidyl 3-(2-pyridyldithio)propionate (SPDP, 0.3 kDa vs. 2.7 kDa, ThermoFisher, UK, #21857). To appraise the percentage labelling the fluorescent signals from the chased samples were calibrated against F-MAL standards. The F-MAL standards[19,37,38] were prepared by mixing ‘fully’ F-MAL and NEM alkylated standards to achieve the desired percentage (e.g., 90% labelled = 9-parts NEM to 1-part FMAL). The fluorescent signals were measured on a plate reader.

*Fig-4G*. To appraise the extent, if any, of inadvertent off-target labelling of other amino acids, such as tyrosine, we added 5 mM 2PB to 75 µg TCEP-reduced NEM-alkylated *X. laevis* egg lysates for 1 h at 24°C. The higher temperature was used to accelerate any side reactions by increasing the likelihood of bimolecular collisions. As a reference, pyridine-2-thione release at 343 nm was monitored in a TCEP-reduced sample in a microplate reader.

*Fig-4H*. To determine how to remove excess 2PB from a sample, we added 5 mM 2PB to 75 µg protein aliquots of NEM-alkylated *X. laevis* oocyte lysates and monitored pyridine-2-thione release at 343 nm after performing the spin column, TCA, and MeOH-based procedures (as above).

### 4.6. Preparing samples for Cleland immunoblotting

After validating the method as above, the samples were lysed in ice-cold buffer (25 mM Tris-HCl, 150 mM NaCl, 1 mM EDTA, 1% NP-40, 5% glycerol, pH 7.1, ThermoFisher, UK, #87787) supplemented with a protease inhibitor tablet (Sigma Aldrich, UK, #11697498001) and 10 mM NEM (ThermoFisher, UK, #23030). The lysates were centrifuged at 5,000 *g* for 5 min at RT to remove insoluble material without solubilising the yolk protein [25]. Reversibly oxidised cysteines were reduced with 5 mM neutral-TCEP (Sigma, UK, #580561) for 30 min. After TCEP was removed, these newly reduced cysteine residues were labelled with 5 mM 2PB for 60 min. All steps were performed in the dark at 4°C with the samples mixed by a 5-10 s vortex every 5 min. Excess NEM or TCEP were removed by size exclusion chromatography with a 6 kDa cut-off spin column (Bio-Rad, UK, #7326222). After determining the protein content via a Bradford assay, excess 2PB was removed via TCA, and the resultant protein pellets were resuspended in non-reducing 4X loading buffer (Bio-Rad, UK, #1610747). A sample boiling step was omitted as it could inadvertently break disulfide bonds, which might lead to undesired cysteine thiyl radicals via homolytic fission of the S-S bond [30].

### 4.7. Streptavidin immunoblotting

To detect the presence (oxidising) or absence (reducing) of the biotin reaction reporter group, 2PB-labelled samples (across a range of inputs 30, 20, 15, 10, and 5 µg protein) were immunoblotted per Cleland immunoblotting with minor modifications. Specifically, the blocked PVDF membranes were incubated with a recombinant streptavidin-conjugated alkaline phosphatase (Abcam, UK, #ab279314) (diluted 1:1000 in TBST containing 5% Superblock™, ThermoFisher, UK, #37580) for 2 h at RT before being washed and imaged. The oxidising and reducing conditions omitted and included DTT, respectively.

### 4.8. Cleland immunoblotting

To implement Cleland immunoblotting, samples containing 20 µg of protein were electrically resolved to their pore size limit on precast 4-15% gradient PAG (Bio-Rad, UK, #4561085) in running buffer (25 mM Tris, 190 mM glycine, 0.1% SDS, pH 8.3) by SDS-PAGE. After carefully excising the gels and rinsing them in ddH_2_O, the gels were soaked in reducing transfer buffer (25 mM Tris, 190 mM glycine, pH 8.3, 0.5% DTT, Sigma, UK #D9779) for 15 min at RT on a plate shaker in a fume hood. The gels were rinsed in ddH_2_O before the proteins were transferred to 0.45 µM low auto-fluorescence PVDF membranes (Bio-Rad, UK, #1620261).

After the transfer step, the PVDF membranes were blocked (1X fluorescent blocking buffer in ddH_2_O, ThermoFisher, UK, #37565) for at least 1 h at RT. The blocked PVDF membranes were immunoprobed with a mouse monoclonal cdc20 antibody raised against the synthetic C-terminal *X. laevis* immunogenic peptide: FEVDPVTKKEKEKARSSKSIIHQSIR [22] (ThermoFisher, UK, #MA1-46040, RRID: AB_1016775). The membranes were incubated in the primary antibody (diluted 1:1000 in TBST containing 5% Superblock™) overnight at RT, washed (3 x 5 min TBST, 1X 10 min ddH_2_0), and probed with an alkaline phosphatase-conjugated mouse-reactive secondary antibody (Abcam, UK, #ab97020) for 1 h at RT, and washed (3 x 5 min).

Impact Vector Red® working solution, per the manufacturer’s instructions (2BScientific, UK, #SK-5100), was applied to the membranes until visible bands appeared. The fluorescent bands were imaged on an Analytik Jena scanner using the appropriate fluorescent excitation (455-495 nm) and emission (607-682 nm) filters in a dark room for 1-10 seconds.

For oxidising Cleland immunoblotting, the method was exactly the same except DTT was omitted from the loading and transfer buffers.

### 4.9. Image processing and analysis

Immunoblot images were analysed on VisionWorks™ software (Analytik Jena, Germany). After subtracting the background, the software automatically calculated the raw densitometry values from the detected bands. The raw data were exported to Microsoft Excel™.

### 4.10. Coding environment

All of the computational analyses were performed using <u>Python</u> in a <u>Google Colaboratory</u> (Colab) coding environment, using the libraries (e.g., pandas) listed in the <u>requirements.txt</u>. The source code is available in the <u>Cleland immunoblotting</u> GitHub repository inclusive of a <u>readme</u> file. The source code is freely available under a <u>MIT license</u> and can be cited using the Zendo <u>DOI</u>.

### 4.11. Computational analyses

To generate the PEG-modified proteins displayed in Figure 1 and 6, the script was <u>Cys_PEG_PDB.py</u> implemented. To generate supplementary data file 1 and 4, modified versions of the script Cys_%25oxI_cal.py were implemented.

For the data in Table 2 and supplementary Figure 2, empirical proteomic data [25] listing the protein ID and estimated copy number (*N*) values for thousands of proteins in *X. laevis* stage XI oocytes were downloaded from (Excel file mmc2 from the supplement). These protein identifiers were matched to UniProt accessions using a customised version of <u>Cys_Expression.py</u> script, available on the <u>Cysteine-</u> <u>i-cloud</u> repository (https://zenodo.org/doi/10.5281/zenodo.13337985). The script used the UniProt API to retrieve the accessions for over 6,000 unique *Xenopus* proteins. For each accession, the number of cysteine residues was determined by parsing the protein sequence data from the UniProt database. This information was compiled into a new Excel sheet, adding a column for the number of cysteine residues. The N data enabled us to calculate the theoretical *i* space (supplementary data file 2).

The number of human proteins that Cleland immunoblotting can measure was estimated by implementing the Cleland_Cys_Proteome.py script on the human reference proteome FASTA file, generating supplementary data file 5.

### 4.12. Constructing the streamlit application

To construct the streamlit application, the <u>Cleland_app.py</u> script was used. For a given input UniProt accession, the source code enables the application to print:

- The number of amino acids.
- The number of cysteine residues.
- The number of cysteine redox proteoforms.
- The number of percentage redox graded bands and their Pascal structure.
- The molecular mass of the 100%-reduced and 100%-oxidised proteoforms (adding 5 kDa per cysteine residue).
- Whether the inputted target is a suitable candidate for Cleland immunoblotting.

The application simulates an immunoblot displaying the estimated location of the percentage redox graded bands (from 100 to 0%-reduced) as an image, that can be downloaded.

### 4.13. Statistical analysis

Data-set normality values were assessed using Shapiro-Wilk and Kolmogorov-Smirnov testing and the experimental design appropriate parametric or non-parametric tests were performed with alpha P ≤

0.05 on <u>GraphPad Prism™ Version 10</u>. When a 1-way ANOVA or non-parametric test was used post-hoc pair-wise comparison tests, such as the Tukey test, were applied as appropriate. All the data are presented as the mean (M) and standard deviation (SD).

### 4.14. Resource

In addition, to the streamlit application and source code, a Cleland immunoblotting protocol file and 2D Cleland immunoblotting file have been uploaded to the Github repository.

## Acknowledgements

The graphical abstract and figure 2 were created using <u>Biorender</u> and exported with a publication permission license. We thank Professor Angus Lamond (Dundee University) and all of the members of the Lamond lab for helpful scientific discussions.

## CRediT Author contributions

**James N. Cobley**: Conceptualisation, methodology, software, formal analysis, investigation, resources, data curation, visualisation, writing-original draft, writing-reviewing and editing. **Anna Noble**: Investigation, resources, writing-reviewing and editing. **Matthew Guille**: Resources, funding acquisition, writing-reviewing and editing.

